# Allosteric communication in Class A β-lactamases occurs via Cooperative Coupling of Loop Dynamics

**DOI:** 10.1101/2020.12.30.424904

**Authors:** Ioannis Galdadas, Shen Qu, Ana Sofia F Oliveria, Edgar Olehnovics, Andrew R Mack, Maria F Mojica, Pratul K Agarwal, Catherine L Tooke, Francesco L Gervasio, James Spencer, Robert A Bonomo, Adrian J Mulholland, Shozeb Haider

## Abstract

Allosteric effects control protein (e.g. enzyme) activity in ways that are not fully understood. Better understanding of allosteric effects, and tools to identify them, would offer promising alternative strategies to inhibitor development. Through a combination of equilibrium and nonequilibrium molecular dynamics simulations, we identify allosteric effects and communication pathways from two distant ligand binding sites to important active site structural elements that control enzymatic activity in two prototypical class A β-lactamases, TEM-1 and KPC-2. Both of these enzymes are important determinants of antibiotic resistance in widespread bacterial pathogens. The simulations show that the allosteric sites are connected to the active site in both enzymes, (e.g. affecting the conformation of the Ω-loop) highlighting how allosteric inhibitors may exert their effects. Nonequilibrium simulations reveal pathways of communication operating over distances of 30 Å or more. In these identified signaling pathways, the propagation of the signal occurs through cooperative coupling of loop dynamics. Notably, 50% or more clinically relevant amino acid substitutions in each enzyme map onto the identified signal transduction pathways. This suggests that clinically important variation may affect, or be driven by, differences in allosteric behavior, providing a mechanism by which amino acid substitutions may affect the relationship between spectrum of activity, catalytic turnover and potential allosteric behavior in this clinically important enzyme family. Simulations of the type presented here will help in identifying and analyzing such differences.

## Introduction

The rise in antimicrobial resistance (AMR) is a growing global public health crisis.^1^ As AMR has continued to spread and many antimicrobial agents have become ineffective against previously susceptible organisms, the World Health Organization recently projected that AMR could result in up to 10 million deaths annually by 2050.^2^ The problem of AMR is particularly urgent given the alarming proliferation of antibiotic resistance in bacteria; pathogens associated with both community-acquired and healthcare-associated infections are increasingly resistant to first-line and even reserve agents.^3^ This not only poses a serious challenge obstacle in fighting common and severe bacterial infections, but also reduces the viability and increases the risks of interventions such as orthopedic surgery and also threatens new antibiotics coming to the market.^4^ AMR risks negating a century of progress in medicine made possible by the ability to effectively treat bacterial infections.

In spite of the advances in the field of antimicrobial chemotherapy, the efficacy, safety, chemical malleability and versatility of β-lactams, makes them the most prescribed class of antibiotics.^5^ Their cumulative use exceeds 65% of all injectable antibiotics in the United States.^6^ β-lactam antibiotics work by inhibiting penicillin-binding proteins (PBPs), a group of enzymes that catalyze transpeptidation and transglycosylation reactions that occur during the bacterial cell wall biosynthesis.^5^ A damaged cell wall results in loss of cell shape, osmotic destabilization, and is detrimental for bacterial survival in a hypertonic and hostile environment.^7^ Of the four primary mechanisms by which bacteria resist β-lactam antibiotics, the most common and important mechanism of resistance in Gram-negative bacteria, including common pathogens such as *Escherichia coli* and *Klebsiella pneumoniae*, is the expression of β-lactamase enzymes.^5^ These enzymes hydrolyze the amide bond in the β-lactam ring, resulting in a product that is incapable of inhibiting PBPs.^8^

The Ambler system of classifying β-lactamase enzymes categorizes them, based on amino acid sequence homology, into classes A, B, C and D.^9,10^ While β-lactamases of classes A, C and D are serine hydrolases, class B enzymes are metalloenzymes that have one or more zinc ions at the active site.^11^ Class A enzymes are the most widely distributed and intensively studied of all β-lactamases.^5^ The hydrolytic mechanism in class A (Figure S1) is initiated by reversible binding of the antibiotic in the active site of the enzyme (formation of the Michaellis complex). This is followed by nucleophilic attack of the catalytic serine (Ser70) on the carbonyl carbon of the β-lactam ring, resulting in a high-energy acylated intermediate that quickly resolves, following protonation of the β-lactam nitrogen and cleavage of the C-N bond, to a lower energy covalent acyl enzyme complex.^12–14^ Next, an activated water molecule attacks the covalent complex, leading to the subsequent hydrolysis of the bond between the β-lactam carbonyl and the serine oxygen, resulting in the regeneration of the active enzyme and release of the inactive β-lactam antibiotic.^5,7,8,12,15–18^

TEM-1 is one of the most common plasmid-encoded β-lactamases in Gram-negative bacteria and is a model class A enzyme.^19^ It has a narrow spectrum of hydrolytic activity that is limited to penicillins and early generation cephalosporins; in contrast, its activity towards large, inflexible, broad-spectrum oxyimino-cephalosporins such as the widely used antibiotic ceftazidime is poor.^8^ However, mutations in the *bla*_TEM-1_ gene have led to amino acid modifications, which allow subsequent TEM-1 variants to hydrolyze broad-spectrum cephalosporins (so-called “extended-spectrum” activity) or to avoid the action of mechanism-based inhibitors such as clavulanate that are used in combination with b-lactams to treat b-lactamase producing organisms.^19^ Another class A enzyme, KPC-2 (*Klebsiella pneumoniae* carbapenemase-2) encoded by the *bla*_KPC-2_ gene is an extremely versatile β-lactamase ^20^ with a broad spectrum of substrates that includes penicillins, cephamycins and, importantly, carbapenems.^20,21^ KPC-2 is also less susecptible to inhibition by clavulanate than is TEM-1. ^22^ Currently, most carbapenem resistance among *Enterobacteriaceae* in the United States and Europe is attributed to plasmid-mediated expression of KPC-type enzymes.^23,24^ Predominant strains of *K*.*pneumoniae* and other *Enterobacterales* continue to be identified as responsible for outbreaks internationally, including the recent first identification of a KPC-2 producing carbapenem-resistant *Klebsiella quasipneumoniae* in Saudi Arabia ^25^. Continued dissemination of KPC makes this one of the β-lactamases of most immediate clinical importance and a key target for inhibitor development.

The structure and activity of Class A β-lactamases have been well studied.^8,28,29^ In spite of sequence differences, class A β-lactamases share the same structural architecture ^30^, as evident from the present 47 structures of TEM-1 and 38 structures of KPC-2, or their engineered variants, deposited in the PDB at the time of this writing. However, despite the wide variety of substrates that TEM-1 and KPC-2 can hydrolyze, their structures are quite rigid. The average mean order parameter, S^2^, as calculated from NMR experiments for TEM-1, is between 0.81-0.94 and almost all class A β-lactamases are conformationally identical. ^31–33^ Loops (e.g. active site loops) play a crucial role in the activity of many enzymes,^34^ including β-lactamases. There is increasing evidence that active site conformations may be influenced by distal loops, connected e.g. through active closure and desolvation, and potentially via networks of coupled motions. ^34,35^ The active sites of TEM-1 and KPC-2 are surrounded by three loops: (a) the Ω loop (residues 172-179), (b) the loop between α_3_ and α_4_ helices, in which a highly conserved aromatic amino acid is present at position 105 and (c) the hinge region, which lies opposite to the Ω loop and contains the α_11_ helix turn (Figure 1, Table S1). Two highly conserved residues, Glu166 and Asn170, which are essential for catalysis, influence the conformation of the Ω loop.^36^ The conformational dynamics of these loops play an important role in enzyme activity, and are probably modulated by evolution.^18,36–40^ For example, we have recently found that differences in the spectrum of activity between KPC-2 and KPC-4 are due to changes in loop behavior. (Tooke et al. in preparation 2020)

**Figure 1:**
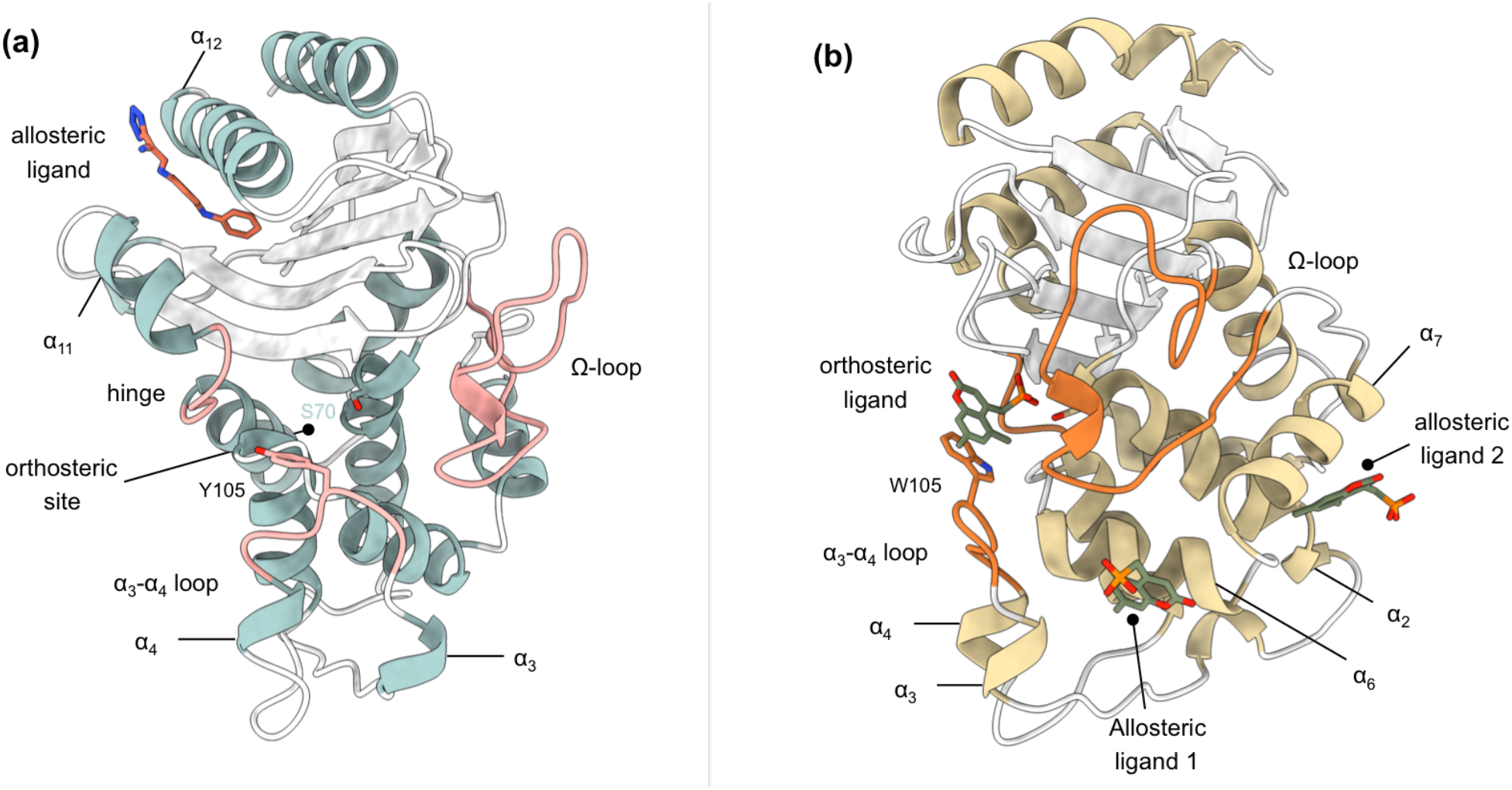
Crystal structures of (a) TEM-1 (PDB id 1PZP)^26^ and (b) KPC-2 (PDB id 6D18)^27^ β-lactamases in complex with ligands bound to allosteric and the orthosteric sites. The helices around the allosteric binding sites and the loops that define the orthosteric binding site are highlighted. In case of KPC-2, allosteric ligand 2 is the site investigated here. See Table S1 for structural nomenclature.

Extensive discussion about the possible contribution of protein dynamics to enzyme catalysis has been advanced.^41–44^ In some enzymes, conformational changes have been identified as necessary for preparing the system for reaction.^34,45^ Several simulation studies, including long timescale and enhanced sampling molecular dynamics (MD) simulations have been reported for TEM-1 and KPC-2 β-lactamases.^12,17,46–48^ MD simulations have explored cryptic pocket formation,^46^ studied protein-ligand interactions,^49^ predicted antibiotic resistance,^12,17,48^ explained the effects of mutations on enzyme specificities,^50^ and investigated conserved hydrophobic networks.^47^ It remains a challenge to directly link conformational heterogeneity and function.

Understanding conformational behavior is relevant to β-lactamase inhibition as well as catalytic mechanism. For organisms producing class A β-lactamases co-administration of susceptible β-lactams with mechanism-based covalent inhibitors (e.g. clavulanate) represents a proven therapeutic strategy and has successfully extended the useful lifetime of penicillins in particular. ^22,51^ However, while the mechanism of direct inhibition by covalently bound inhibitors is well established,^22^, the possibility of exploiting sites remote from the active center in allosteric inhibition strategies is less explored, and where this has been achieved ^26,27,48^ the structural changes occurring as a result of ligands binding or unbinding to allosteric sites, and the relay of structural communication that leads to inhibition are not well understood. The conformational rearrangements that take place upon ligand (un)binding in allosteric sites and their potential connection to the β-lactamase active site, are the focus of this study.

Here, we employ a combination of equilibrium and nonequilibrium molecular dynamics (MD) simulations to identify and study the response of two class A β-lactamases, namely TEM-1 and KPC-2, to the (un)binding of ligands at sites distant from the active site. Nonequilibrium simulations applying the Kubo-Onsager approach ^52,53^ are emerging as an effective way to characterize conformational changes and communication networks in proteins ^54–57^.

To the best of our knowledge, this is the first application of this nonequilibrium MD approach to study enzymes including β-lactamases, whose ultrafast turnover rates can approach the diffusion limits for natural substrates (∼10^7^-10^8^ M^-1^s^-1^).^15^ We perform 10 µs of equilibrium MD simulations of TEM-1 and KPC-2, with and without ligands present in their allosteric binding sites. These simulations identify conformational changes in the highly dynamic loops that shape the active site and structurally characterize the dynamics of the formation and dissolution of the allosteric pocket. We also carry out an extensive complementary set of 1600 short nonequilibrium MD simulations (a total of 8 µs of accumulated time), which reveal the response of the enzyme to perturbation and identify pathways in the enzymes that connect the allosteric site to other parts of the protein. These simulations demonstrate direct communication between the allosteric sites and the active site. The results show that this combination of equilibrium and nonequilibrium MD simulations offers a powerful tool and a promising approach to identify allosteric communication networks in enzymes.

## Results and Discussion

### Equilibrium Simulations of Apo_EQ_ and IB_EQ_ states

To explore the conformational space of TEM-1 and KPC-2 in the Apo_EQ_ (no ligand) and IB_EQ_ (inhibitor bound) states, we started by running a set of equilibrium simulations (20 replicas of 250 ns each) that resulted in 5 µs of accumulated simulation time per system. Conformational changes during the simulations were assessed using their Cα root mean-squared deviation (RMSD) profiles (Figure S3). The simulated systems were considered equilibrated beyond 50 ns as shown by RMSD convergence. In each case, the proteins remained close to their initial conformation during the course of 250 ns (Figure S3a). The average RMSD for Apo_EQ_ and IB_EQ_ states were between 0.10-0.12 nm for all systems (Table S2). The low RMSD values are consistent with previously published results, which have also shown class A β-lactamase enzymes to be largely rigid and conformationally stable when studied on long time scales and rarely divergent from the initial structure. ^31,47^ Conventional RMSD fitting procedure using all Cα atoms failed to separate regions of high versus low mobility. To resolve such regions, we used a fraction (**%**) of the Cα atoms for the alignment. Beyond this fraction, there is a sharp increase in the RMSD value for the remainder of the Cα atoms (Figure S3b). At 80%, the core of TEM-1 could be superimposed to less than 0.064 nm and 0.074 nm for Apo_EQ_ and IB_EQ_ states, respectively (Figure S3bi).

In the KPC-2 Apo_EQ_ state, the RMSD of 80% of the Cα atoms was below 0.060 nm, while the same subset of atoms had an RMSD below 0.066 nm in the IB_EQ_ state (Figure S3bii). This 80% fraction of Cα atoms constitutes the core of the enzyme and did not show any divergence from the initial reference structure (Figure S3c). RMSD values for the remaining 20% of Cα atoms that diverged from the core varied between 0.16 to 0.227 nm. This apparent rigidity is consistent with the experimental finding, based upon e.g. thermal melting experiments ^58^, that KPC-2 is a more stable enzyme than many other class A β-lactamases such as TEM-1. Some large conformational changes were observed in all replicates; these involved changes in conformations of the loops that connect secondary structural elements (Figure S4). To further validate the stability of the two systems, we analyzed structural properties including the radius of gyration (Rg; Figure S5), solvent accessible surface area (SASA; Figure S6) and the secondary structure of each enzyme over the simulated time (Figure S7). The values for these properties are listed in Table S2.

### Conformational Sampling of Equilibrated Apo_EQ_ and IB_EQ_ states

The convergence of the dynamical behavior of the systems in the equilibrated trajectories was assessed using Principal Component Analysis (PCA) (Figure S8).^59,60^ For the PCA of the equilibrium simulations, all 20 replicates from Apo_EQ_ and IB_EQ_ were combined before the analysis. In TEM-1, 101 eigenvectors were required to describe 90% of the variance. Similarly, 103 eigenvectors described 90% of the variance in KPC-2. PCA was able to extract and filter dominant motions from the sampled conformations and define their respective essential space. In both TEM-1 and KPC-2, these motions relate to the hinge-α_11_, α_12_, α_11_-β_7_, β_8_-β_9_, β_1_-β_2_, α_9_-α_10_, α_2_-β_4_, α_7_-α_8_, α_1_-β_1_, α_12_-β_9_, α_1_-β_1_, α_3_-turn-α_4_ and Ω loop (Figure S9 and S10). We next made a pairwise comparison between the Apo_EQ_ and IB_EQ_ states by calculating the dot product matrix between the eigenvectors found from PCA in the Apo_EQ_ protein, with those found from PCA in the IB_EQ_ state. Such comparison allows quantitative assessment of similarity between the dynamics in the two different systems. TEM-1 Apo_EQ_ and IB_EQ_ simulations have a subspace overlap of 74% and an average maximum dot product of 0.50. The most significant similarity observed between TEM-1 Apo_EQ_ and IB_EQ_ is in PC2, with an inner product of 0.83 (Figure S8e). This PC represents the motions in the hinge-α_11_, α_12_, α_12_-β_9_, and α_1_-β_1_. Similarly, the KPC-2 simulations have a subspace overlap of 76% and an average maximum dot product of 0.49. PC2 from Apo_EQ_ and PC1 from IB_EQ_ display the largest similarity (Figure S8f). The motions observed in these PCs are dominated by the loops α_7_-α_8_ and the α_3_-turn-α_4_ helices (Figure S10). The overall eigenvectors of the Apo_EQ_ and IB_EQ_ simulations, in both systems, define a subspace that exhibits a conserved dynamic behavior between TEM-1 and KPC-2. In TEM-1 and KPC-2 IB_EQ_ states, the unrestrained ligands remained bound in the allosteric sites throughout the course of the simulations, and despite fluctuations, both systems displayed similar dynamics. The PCA analyses suggest that, for each enzyme, the Apo_EQ_ and IB_EQ_ simulations occupy the same conformational (essential dynamics) subspace, though not always sampling the same regions in that space.

### Ligand-induced Structural and Dynamical Changes

A ligand that binds to an allosteric site can control protein function by affecting the active site.^61^ This generally occurs by altering the conformational ensemble that the protein adopts.^61,62^ To probe how ligand binding to an allosteric site affects the dynamics of β-lactamases, we calculated the Cα root mean-square fluctuation (RMSF) for both Apo_EQ_ and IB_EQ_ states. Higher RMSF values correspond to greater flexibility during the simulation. Although the Cα RMSF profiles for Apo_EQ_ and IB_EQ_ states are similar, indicating similar dynamics, there are some discernible differences (Figure 2).

**Figure 2:**
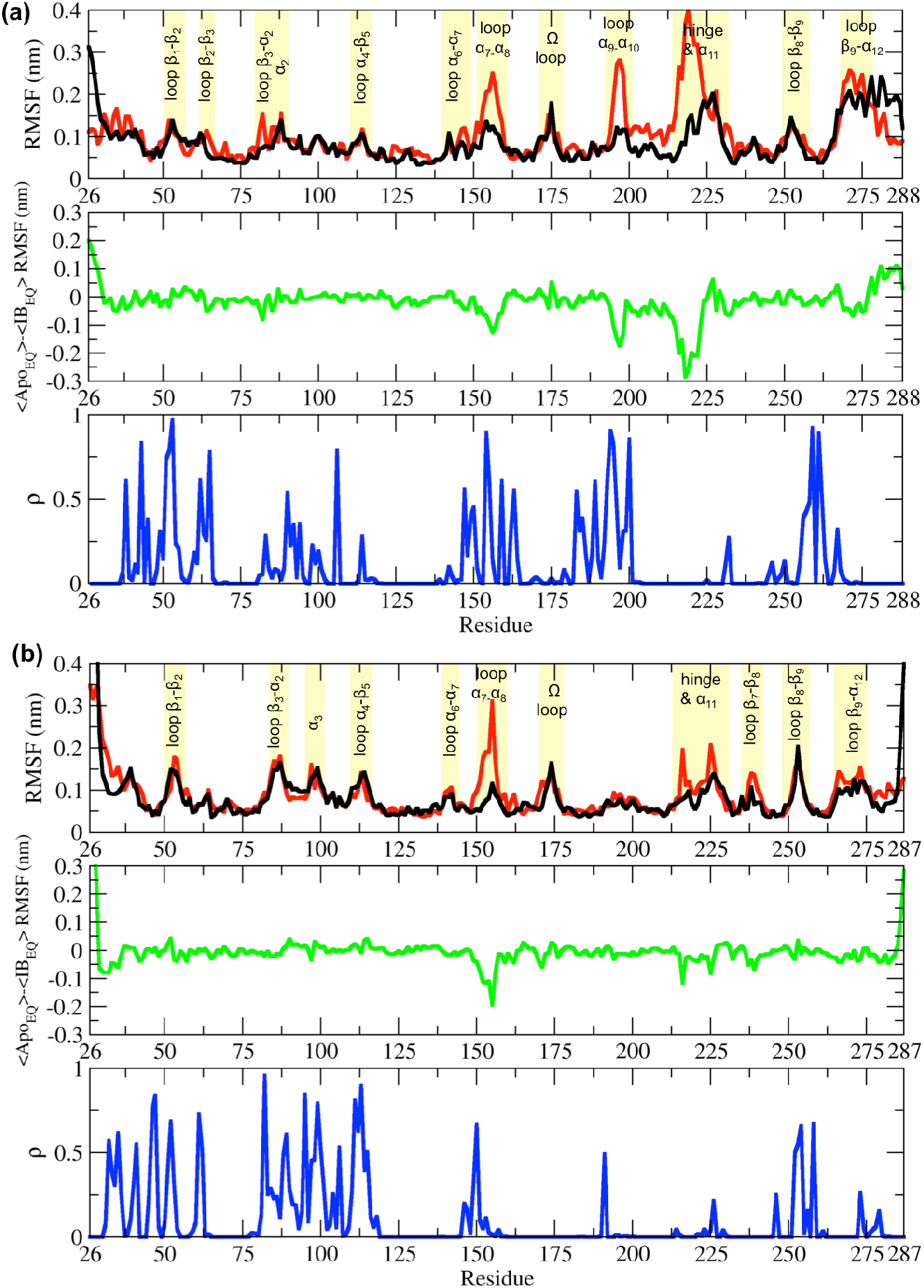
RMSF differences between the Apo_EQ_ and IB_EQ_ states of the (a) TEM-1 and (b) KPC-2 systems. The average change in RMSF in the Apo_EQ_ (black), the IB_EQ_ (red), the difference Apo_EQ_-IB_EQ_ (green), and the associated ρ value (blue) is illustrated. The ρ values were obtained by conducting a Student’s t-test to compare Apo_EQ_ and IB_EQ_ systems and to assess the significance of the differences.

In simulations of TEM-1 and KPC-2, the hydrophobic core of the enzyme is stable and shows limited fluctuations. Most of the RMSF variance is observed in loops that connect secondary structural elements (Figure 2). In TEM-1 IB_EQ_, higher fluctuations are observed predominantly in three distinct regions when compared with the Apo_EQ_ enzyme; in the loops between helices α_7_ and α_8_ (residues 155-165), α_9_ and α_10_ (residues 196-200) and the hinge region including α_11_ helix (residues 213-224) (Figure 2A). The α_11_ and the α_12_ helices are part of a highly hydrophobic region that also constricts the allosteric pocket in all TEM-1 Apo crystal structures. Binding of the ligand disrupts the hydrophobic interactions within this region, resulting in the opening of the allosteric pocket between α_11_ and α_12_ helices.^26^

It should be noted that the starting Apo_EQ_ structure of TEM-1 was generated from the IB crystal structure, by the removal of the ligand from the allosteric binding site. During the Apo_EQ_ simulations, α_12_ helix behaves like a lid and closes over the empty, hydrophobic, allosteric binding site, and thus displays high RMSF at the C-terminal end of the enzyme. This conformational change recovers the structure of the Apo crystal form, as observed e.g. in PDB id 1ZG4 ^63^, to an overall superimposable RMSD of ∼0.07 nm. A similar motion of an a-helix has also been observed previously for a major urinary protein, where the binding cavity is so hydrophobic that the water molecules prefer to avoid it, even in the absence of the ligand.^64^ The rest of the loops displayed comparable fluctuations in both Apo_EQ_ and IB_EQ_ states.

The differences between the Apo_EQ_ and IB_EQ_ states were of similar magnitude in KPC-2. In KPC-2 IB_EQ_, more extensive fluctuations than in Apo_EQ_ were also observed in the loops between α_7_-α_8_ (residues 156-166), the hinge region, around α_11_ (residues 214-225) and in the loop between β_7_-β_8_ (residues 238-243) (Figure 2b). Conversely, fluctuations are slightly higher in the Apo_EQ_ than IB_EQ_ state in the loop leading into the Ω loop from α_7_ helix (residues 156-166). Overall, however, RMS fluctuations are similar in analogous regions of the IB_EQ_ and Apo_EQ_ states in both TEM-1 and KPC-2, highlighting the conservation of structural dynamics in class A β-lactamases. However, there were some fluctuations that were unique and limited to each enzyme (Figure 2).

In both TEM-1 and KPC-2 IB_EQ_ states, interactions of the ligands in their respective allosteric binding sites contribute to enhanced fluctuations (i.e. larger than in the Apo forms) of the local structural elements (Figure S11). The sites in which the ligands bind are very different. In TEM-1, the binding site is deep and forms a hydrophobic cleft. The ligand penetrates to the core of the enzyme and is sandwiched between α_11_ and α_12_ helices. (Horn and Shoichet 2004) The FTA ligand remains tightly bound in the allosteric pocket throughout the simulations (Figure S12).

In KPC-2, the allosteric binding site is shallow and solvent-exposed even in the absence of the ligand. Although the distal end of the pocket is hydrophobic, there are some polar amino acids on the proximal surface (e.g. Arg83 and Gln86), which are exposed to the solvent. This shallow site forms a part of a larger pocket that is occluded by the side chain of Arg83 (α_7_ helix). In some of our IB_EQ_ simulations, the Arg83 side chain rotates, leading to the opening of a larger hidden pocket. This enlarged space is now accessible to the ligand for exploring various interactions. The tumbling of GTV increases the fluctuations in the complex (Figure S11c,d), however, the ligand does not leave the binding site (Figure S12).

To further highlight the structural changes occurring as a result of ligand binding, positional Cα deviations were calculated between IB_EQ_ and Apo_EQ_ systems for the equilibrated part of the simulations (Figure 3a,b). The Cα deviation values plotted are an average between simulation taken by combining all trajectories from Apo_EQ_ and IB_EQ_ simulations for that particular system. This is one of the simplest approaches, which can determine residues undergoing largest structural rearrangements. The averaged Cα positional deviations are mapped onto the averaged Apo_EQ_ structure to visualize the largest relative displacements in three-dimensions (Figure 3c,d).

**Figure 3:**
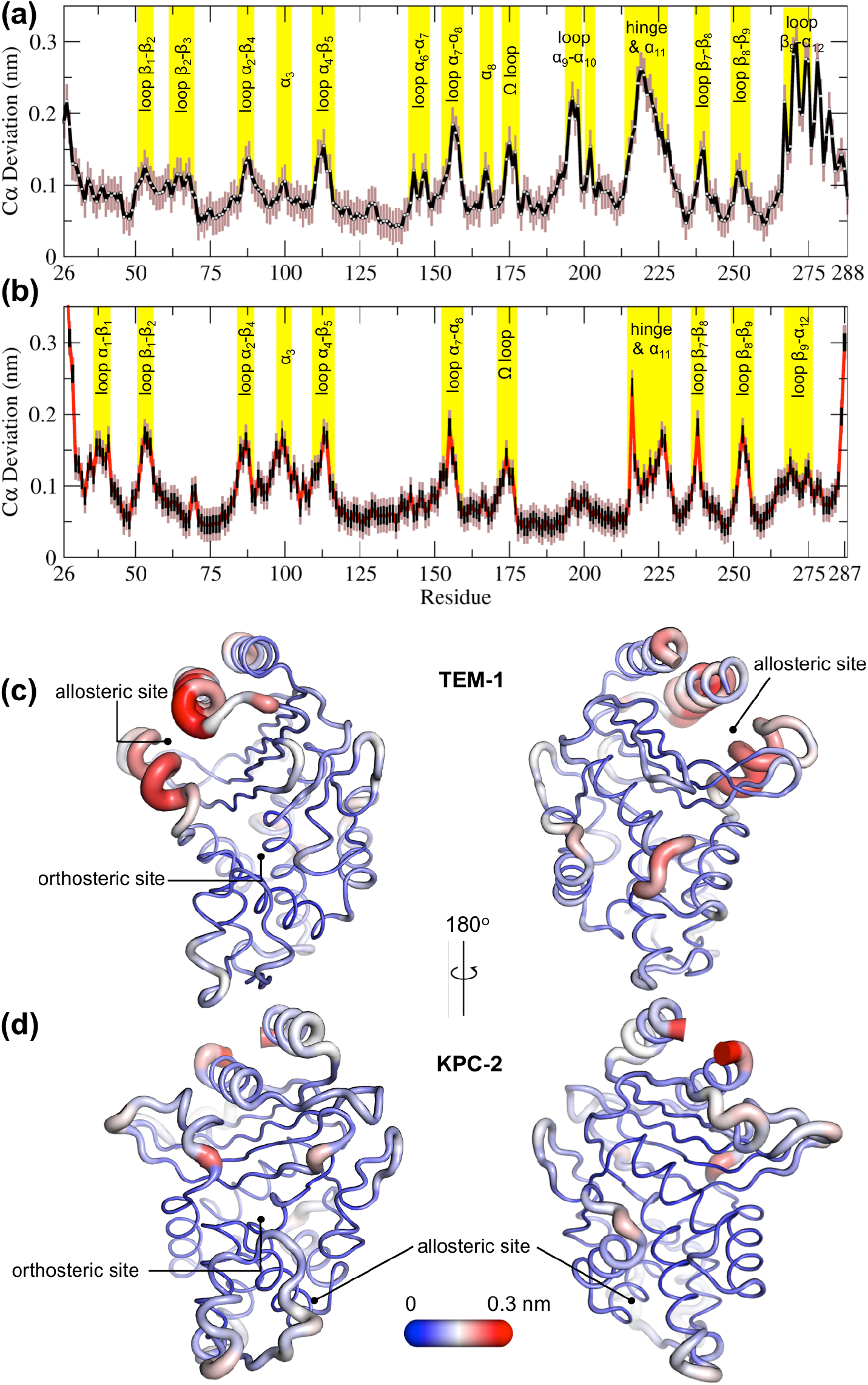
Average positional Cα deviations between the Apo_EQ_ and IB_EQ_ states of (a) TEM-1 and (b) KPC-2. Important structural motifs are highlighted and labeled on the plots. The brown vertical lines represent the standard deviation of the mean. The averaged Cα positional deviations mapped on the averaged Apo_EQ_ structures of (c) TEM-1 and (d) KPC-2, to visualize the largest relative displacements. The average deviation was determined from a combination of all 20 Apo_EQ_ and 20 IB_EQ_ trajectories. The thickness of the cartoon corresponds to the Cα deviation.

The hydrophobic cores of both TEM-1 and KPC-2 β-lactamase enzymes show little or no conformational change. The major differences between the Apo_EQ_ and IB_EQ_ states are in the loops connecting different secondary structure elements. In TEM-1, Cα deviations are observed in the loops between α_4_-β_5_ (residues 112-116), α_7_-α_8_ (residues 155-166), Ω loop (residue 172-179), α_9_-α_10_ (residue 196-200), hinge and α_11_ (residues 213-224), β_7_-β_8_ (residues 238-243) and β_9_-α_12_ (residues 267-272). There are some relatively minor deviations observed in loops β_1_-β_2_ (residues 51-55), β_2_-β_3_ (residues 61-65), α_2_-β_4_ (residue 86-93), α_6_-α_7_ (residues 143-144), β_8_-β_9_ (residues 252-258) and at the pivot of α_3_ helix (residues 98-101). The hinge region and residues in helices α_11_ and α_12_ display the largest deviations. This is also in agreement with other experimental data where the connection existing between the active site and the allosteric pocket studied in TEM-1 in the presence of BLIP inhibitor, seems to be mostly due to hinge region motions.^65^

The structural dynamics observed in KPC-2 were slightly different from TEM-1. In KPC-2, prominent Cα deviations were observed in the loops between β_1_-β_2_ (residues 51-55), α_2_-β_4_ (residue 88-93), α_4_-β_5_ (residues 114-116), α_7_-α_8_ (residues 156-166), Ω loop (residue 172-179), β_7_-β_8_ (residues 238-243), β_8_-β_9_ (residues 252-258), in the loop between β_4_-α_3_ leading up to the proximal end of α_3_ (residues 94-102) and in the hinge/α_11_ helix (residues 214-225). There are some minor deviations observed in α_1_-β_1_ (residues 39-42) and β_9_-α_12_ (residues 266-270). The most important ligand-induced Cα deviation is observed in the loop connecting α_4_ helix to β_5_ strand (residues 114-116). The deviation of the α_4_-β_5_ loop together with the deviation observed in the loop between β_4_-α_3_ leading into α_3_ helix (residues 96-102) has the potential to deform the α_3_ helix-turn-α_4_ helix. The

β_4_-α_3_ and α_4_-β_5_ loops form the basal pivot joint of the α_3_ and α_4_ helices and maintain the correct positioning of this helix-turn-helix at the periphery of the enzyme active site. The correct positioning of this loop is important as Trp105 lies on this loop. Mutagenesis studies have shown that a highly conserved aromatic amino acid at position 105 in class A β lactamases (Tyr105 in TEM-1, Trp105 in KPC-2) is located at the perimeter of the active site and plays a crucial role in ligand recognition via favorable stacking interactions with the β-lactam ring.^66,67^ The aromatic side chain at position 105 coordinates the binding of substrates not only via stacking and edge-to-face interactions but by also adopting “flipped-in” or “flipped-out” conformations.^47,66,68^ This has been proposed based on the conformations observed in the available crystal structures and confirmed by enhanced sampling molecular dynamics simulations.^47,69^ Any perturbation that alters the conformation of α3-turn-α4 helix or deforms the α_3_-α_4_ pivot region would prevent α_3_ and α_4_ helices from correctly shaping the active site of the enzyme. This would result in the aromatic residue at 105 partially detaching from the edge of the active site and being unable to stabilize the incoming substrate as required for efficient catalysis. This explains the loss of b-lactam resistance in strains expressing KPC variants at positions 102 or 108, as established in the MIC experiments reported previously.^47^

### Signal propagation from the allosteric site

To study signal propagation from the two allosteric sites, we ran 800 short nonequilibrium (NE) simulations, with a total sampling time of 4 μs for each system. The nonequilibrium simulations were initiated from regular intervals of the equilibrated part of the long IB_EQ_ simulation, starting at the 50 ns time point (Figure S2). In each simulation, the ligand was removed from its binding site and the resulting system was further simulated for 5 ns. The response of the system to the perturbation was determined using the Kubo-Onsager approach developed by Ciccotti *et al*.^52,53,70^. In this approach, the time evolution of the conformational changes induced by ligand removal can be determined by comparing the Apo_NE_ and IB_EQ_ simulations at equivalent points in time. The subtraction method cancels the noise arising from fluctuations of the systems and allows residues that are involved in signal propagation to be identified. The disappearance of the ligand from its binding site generates a localized vacuum, against which there is an immediate structural response by the enzyme. As the simulation progresses, the cascading conformational changes in response to the perturbation (removal of ligand) show the route by which structural response is transmitted through the protein.

This approach has identified a general mechanism of signal propagation in nicotinic acetylcholine receptors, by analyzing their response to deletion of nicotine.^56^ The difference in the position of Cα atoms is calculated between the short Apo_NE_ and IB_EQ_ simulations at specific time points. These differences are then averaged over all pairs of simulations to reveal the structural conformations associated with this response (Figure 4) and their statistical significance. The Cα coordinates of each residue in the Apo_NE_ were subtracted from the corresponding Cα atom coordinates of the IB_EQ_ simulation at specific points in time, namely 0.05, 0.5, 1, 3, and 5 ns. This resulted in a difference trajectory for each pair of simulations. The difference trajectories are averaged over the set of 800 simulations for each system. The low standard error calculated for the average between the Apo_NE_ and IB_EQ_ demonstrates the statistical significance of the results. Due to the short timescale (5 ns) of the nonequilibrium simulations, only small amplitude conformational changes will be observed.

**Figure 4:**
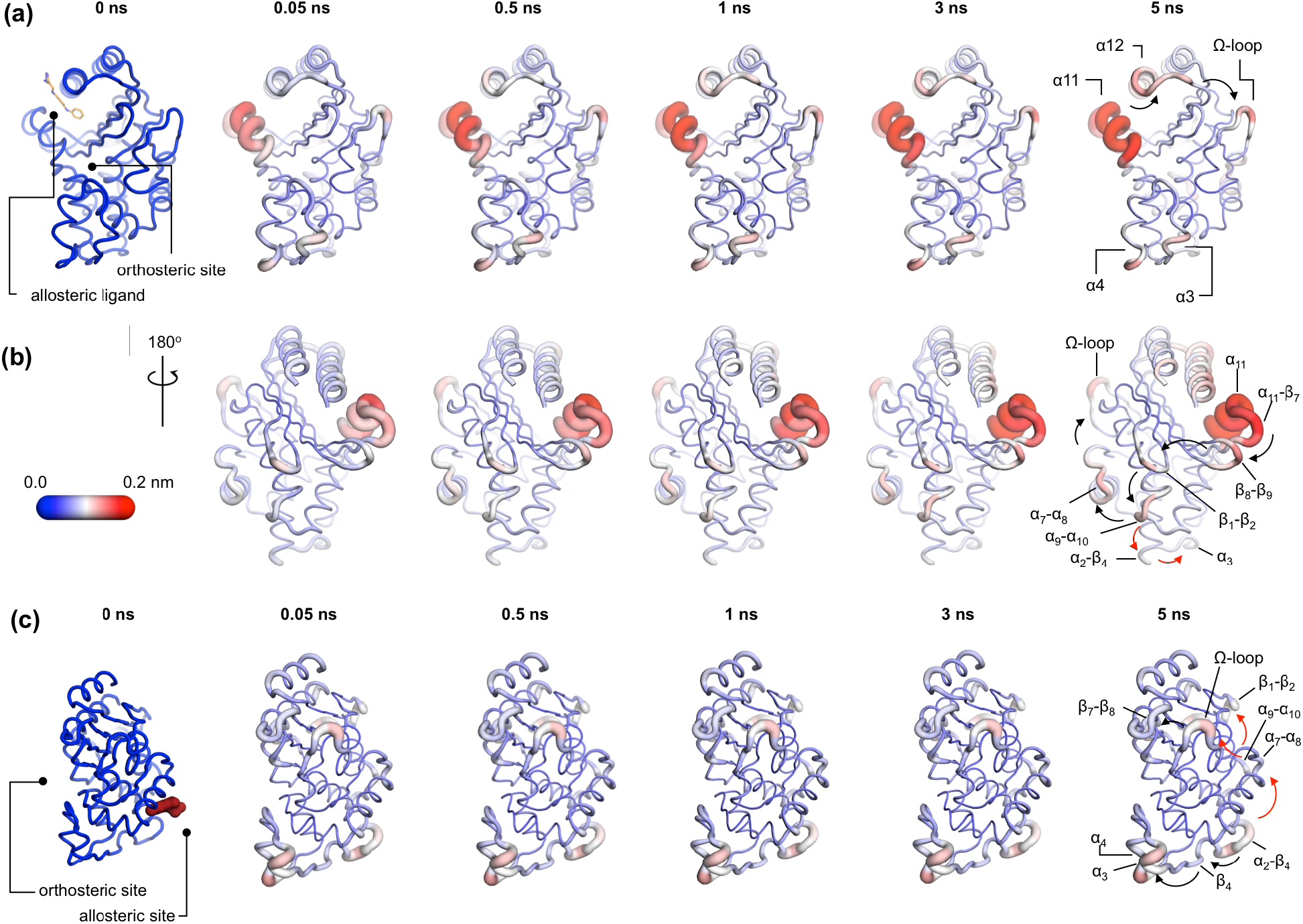
Communication pathways in (a, b) TEM-1 and (c) KPC-2. The average Cα deviations correspond to the average difference in the position of each Cα atom between all 800 pairs of IB_EQ_ and Apo_NE_ simulations at specific time points. The averaged Cα deviations are mapped on the average Apo_EQ_ structure. The arrows mark the direction of the propagation of the signal, caused by the perturbation (removal of the ligand). The red and the black arrows highlight different paths taken by the propagating signals (Also see movies S1, S2 and S3)

In TEM-1, the allosteric site is sandwiched between the α_11_ and α_12_ helices. Adjacent to this binding site is the hinge region (residues 213-218), whose dynamics have previously been examined by NMR and shown to have low order parameters indicating high mobility.^31,33^ This is also the site of perturbation in the nonequilibrium simulations and so the point of origin of the allosteric signal. Located on the loop between the distal end of the α_11_ helix and β_7_ is a highly conserved Trp229 residue. The indole ring of Trp229 is sandwiched between two other highly conserved residues, Pro226 and Pro251 present in loops α_11_-β_7_ and β_8_-β_9_ respectively. The π/aliphatic stacked arrangement of tryptophan-proline is a very tight interaction and is similar in geometry to that observed in complexes of proline-rich motif-binding families, including the EVH1 and GYF binding domains, with their peptide ligands.^71–74^ The perturbation destabilizes this stacked arrangement resulting in an extension of an inherently highly mobile region. After 50 ps of simulation, the Cα deviations have propagated and can be observed in the loop between β_1_ and β_2_. Interestingly, the loops at the basal pivot of α_3_ and α_4_ also responded rapidly to ligand removal. These loops are ∼33 Å away from the allosteric binding site and can affect the spatial position of the turn between helix α_3_ and α_4_. The α_3_-turn-α_4_ helix forms the boundary of the active site, and it is on this turn where the Tyr105 residue, important for substrate recognition is positioned. These results clearly demonstrate the coupling between the distal allosteric site and catalytically relevant regions of the enzyme. As the signal propagates within the protein, there is a gradual and cumulative increase in the Cα deviations in the aforementioned loops. In particular, the loop between the α_9_ and α_10_ helices, which is positioned just below the β_1_-β_2_ loop, displays high deviations and forms a focal point for the signal to bifurcate in two directions. First, major deviations are observed laterally towards loop α_7_-α_8_ and onwards into the Ω loop (Figure 4a,b). Second, more minor deviations move into the loop between α_2_-β_4_ and onwards into the basal pivot of α_3_-turn-α_4_ helix. There is another shorter route at the top of the enzyme that the signal can take to go from the allosteric binding site to the Ω loop, via the proximal end of α_12_ helix and across the loop between β_9_-α_12_ helix (Figure 4a,b).

In KPC-2, the allosteric pocket is shallower and lies between helices α_2_ and α_7_. Residues from three loops (α_6_-α_7_, α_7_-α_8_ and α_2_-β_4_) are in close proximity to this binding site. An additional loop, α_9_-α_10_, is linked to this binding site via the distal end of the α_2_ helix. The perturbation in this binding site results in enhanced mobility of the α_2_-β_4_ loop, which leads directly into β_4_ and onwards to the basal pivot of the α_3_ helix. The proximal end of the α_3_ helix and the distal end of the α_4_ helix, which forms the pivot point of the α_3_-turn-α_4_ structure, display high deviations (Figure 4c). The highly conserved aromatic amino acid, Trp105, is located on this turn. The distance between the allosteric binding site and the α_3_ helix is ∼27 Å. Other major deviations are also observed in the Ω loop as the simulation progresses (Figure 4c). The Ω loop is directly linked to the allosteric binding site via loop α_7_-α_8_. Some minor deviations are also observed in the loop connecting β_9_ and the α_12_ helix.

In both TEM-1 and KPC-2, the removal of the ligand at the beginning of the nonequilibrium simulations does not result in large conformational changes. The subsequent Cα deviations trace the route of the propagating signals (Figure S12). In TEM-1, α_11_ and the hinge region, loop β_1_-β_2_ and loop β_8_-β_9_ respond rapidly to the perturbation and display comparable RMSD values to the equilibrated simulations. Similarly, in KPC-2, only loops α_2_-β_4_ and α_7_-α_8_ respond rapidly to the perturbation. The rest of the structural elements take longer to respond, and their conformational rearrangements are not fully sampled in the Apo_NE_ simulations. It is worth emphasizing that while the short nonequilibrium simulation can be an excellent tool to study an immediate structural response towards a perturbation, the timescale of nonequilibrium MD does not represent a real timescale and thus should not be compared directly with equilibrium simulations. Nevertheless, nonequilibrium MD can identify the sequence of events and pathways involved.

The perturbations of the two enzymes here are different but show some striking common features. In both TEM-1 and KPC-2 systems, even though the point of origin of perturbation (i.e. allosteric site) is different, the signal leads to common endpoints at the pivot of α_3_-turn-α_4_ helix and in the Ω loop. Thus, simulations of two disparate class A b-lactamases, starting from two distinct allosteric sites, identify a common mechanism by which catalytic activity may be disrupted through conformational changes close to the active site. The results from the nonequilibrium simulations also correlate well with experimental data, which suggest that the Ω loop plays a critical role in ligand binding by altering the conformation of Glu166 and Asn170 which are involved in both acylation and deacylation reactions.^12,18,19,22,36^

### Dynamic cross-correlation analysis of surface loops

Dynamical cross-correlation analysis provides information about the pathways of signal propagation and also some insights into the timescales of allosteric communication in TEM-1 and KPC-2 β-lactamases. The dynamic cross-correlation maps (DCCM) have been previously used to identify networks of coupled residues in several enzymes by us. ^75–77^

Using a similar approach, Dynamic Cross-Correlation Maps (DCCMs) were calculated for the Apo_EQ_ and IB_EQ_ simulations and also for the Apo_NE_ nonequilibrium simulations (Figure 5). In these figures, the green regions represent no to slightly positive correlations, while yellow regions represent moderate negative correlations. Negative correlations imply residues moving towards or away from each other in correlated fashion (such as shown by fluctuating hydrogen bonds); for large regions this represents global conformational fluctuations (also referred to as breathing motions).^75^ The results depicted in Figure 5a indicate that in the case of TEM-1 Apo_EQ_ (Figure 5a, left), α_11_ helix shows high negative correlation with α_12_ terminal helix. This represents the lid motion of α_12_ helix, which moves to shut the empty, hydrophobic, allosteric binding site in the TEM-1 Apo_EQ_ structure (see above). This motion is, however, not observed in the ligand bound TEM-1 IB_EQ_ simulations. The TEM-1 IB_EQ_ system shows a substantial increase in correlations, representing changes in the dynamical communications due to the presence of the allosteric ligand (Figure 5a, middle). The binding of the ligand changes the overall global conformational fluctuations of TEM-1, as represented by the increase in yellow regions in the DCCMs. Furthermore, a number of negative correlations (encircled red regions in DCCMs) also increase in other regions of the protein on ligand binding. The DCCM collectively computed from all nonequilibrium trajectories for TEM-1 (Figure 5a, right, Figure S14) also shows a further increase in the areas of negative correlations (encircled). Interestingly, DCCM also identifies the pathway of allosteric communication (Figure S14), with notable correlations between the regions β_1_-β_2_:α_2_-β_4_, α_3_-α_4_:α_2_β_4_, β_4_-α_3_:α_7_-α_8_, β_3_-α_2_:Ω, α_9_-α_10_:β_1_-β_2_, β_3_-α_2_:β_8_-β_9_, α_5_-α_6_:α_12_, hinge-α_11_:α_1_-β_1_, β_8_-β_9_:α_4_-β_5,_ β_7_:α_12_ and α_11_:α_12_. These results indicate that the presence of ligand in TEM-1 increases the dynamic communication between regions that are independent in the Apo_EQ_ simulations. This is particularly evident in the nonequilibrium trajectories that show the largest changes from the case of Apo_EQ_ TEM-1, identifying changes in correlation as the system adjusts to the absence of the ligand.

**Figure 5:**
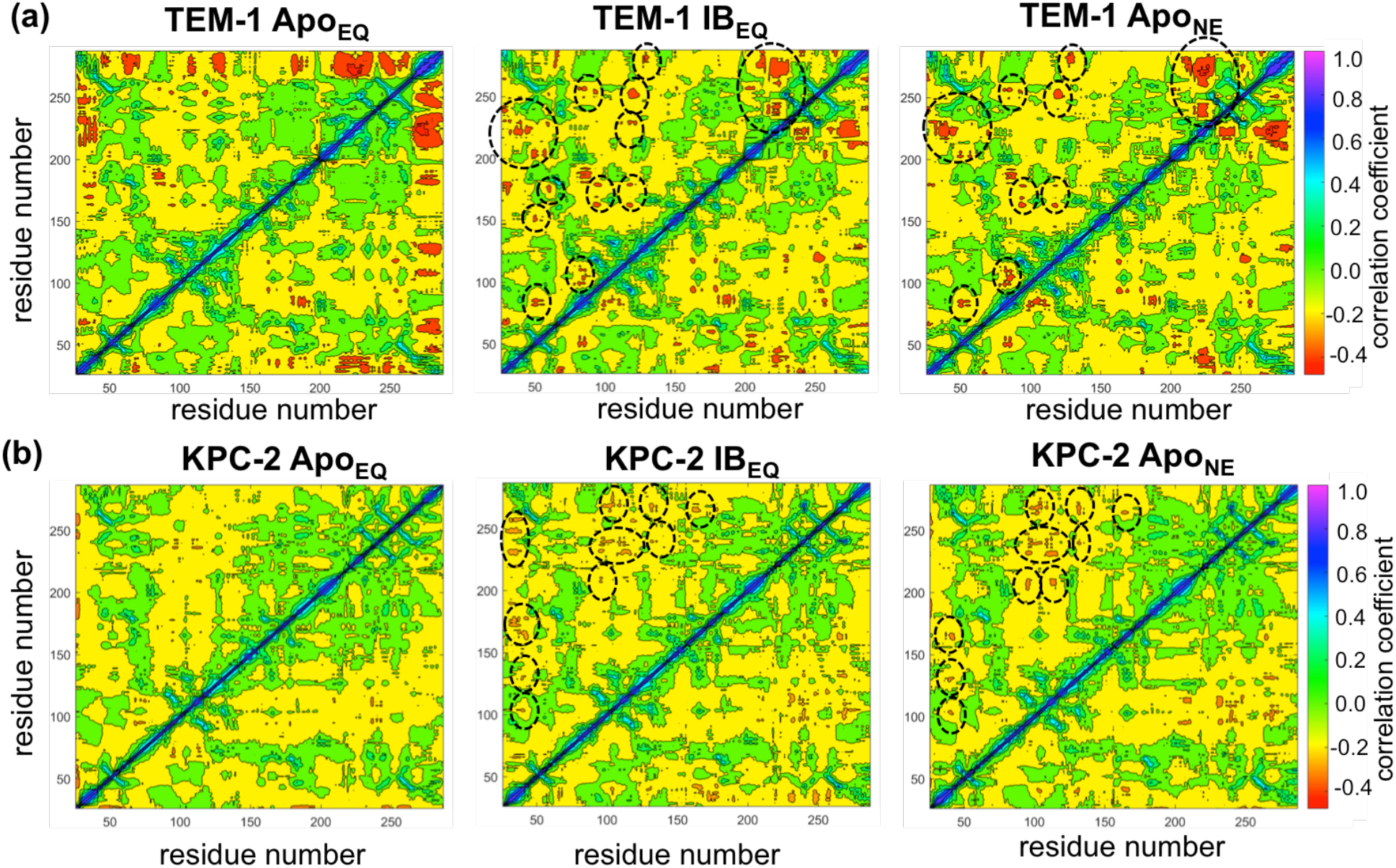
Dynamic Cross-Correlation Maps (DCCMs) computed for (a) TEM-1 and (b) KPC-2 Apo equilibrium (Apo_EQ_), inhibitor-bound equilibrium (IB_EQ_) and Apo nonequilibrium (Apo_NE_) trajectories. The DCCMs for equilibrium trajectories were calculated as an average of 20 replica simulations, while the Apo nonequilibrium DCCM indicates an averaged DCCM from an ensemble of 40 short (5 ns) MD trajectories. Green regions indicate no correlation; yellow indicates moderate negative correlation while orange and red indicate significant negative correlations, while blue regions indicate positive correlations. In TEM-1 Apo_NE_, regions showing significant changes from Apo_EQ_ and IB_EQ_ bound simulations have been marked by black dashed ellipses.

KPC-2 shows even more interesting behavior (Figure 5b). Simulations of Apo_EQ_ KPC-2 show overall more correlated regions than TEM-1 Apo_EQ_ system (as indicated by the more extensive yellow regions in the DCCM), with further increases in the presence of the inhibitor (indicated by a number of orange regions). However, the DCCM collectively computed from all nonequilibrium trajectories for KPC-2 shows a reduction in regions of cross-correlations; a contrast from the case of TEM-1. To obtain a better understanding, the DCCMs from individual 5 ns nonequilibrium trajectories were also computed and analyzed. These reveal interesting trends as depicted in Figure S15. For most nonequilibrium trajectories, the maps are similar with a decrease in dynamic correlations, however, for several trajectories (shown in Figure S15) the maps indicate a significant increase in the correlations. The DCCMs computed from individual trajectories show behavior similar to averaged nonequilibrium trajectories in TEM-1 with a number of regions showing high negative correlations (as highlighted by wide-spread presence of small red regions in the DCCMs). Overall, these results indicate that the perturbation in KPC-2 generates a dynamical response that is much faster than that observed in TEM-1. A plausible explanation for the faster response in KPC-2 is that the more solvent-exposed ligand binding site is surrounded by dynamic surface loops that respond to the perturbation more quickly than the allosteric binding site in TEM-1, which is buried in the hydrophobic core of the protein. This is consistent with the experimental observations that motions can occur on different timescales and can vary greatly between different β-lactamases.^31^

### Relating enzyme dynamics to positions of substitution in TEM-1 and KPC-2 clinical variants

A number of clinical variants that extend hydrolytic activity to encompass additional b-lactams such as oxyiminocephalosporins, and/or enhance enzyme stability, have been identified for both the TEM-1 and KPC-2 β-lactamase enzymes.^8,78,79^ Some of these have been crystallized and their protein structure deposited in the PDB. While many of these amino acid substitutions (for example TEM-1 mutations at residues Glu104 in the α_3_-turn-α_4_, Arg164 on the Ω-loop and Ala237, Gly238 and Glu240 on the β_7_ strand) directly affect important structural features such as the active site or the Ω loop, some are of uncertain structural significance. Even when enzyme structures are known, the connections between the positions of clinical variants, protein structure and their functional implications are often not clear. There is particular uncertainty and interest in the effects of mutations more distant from the active site.

To assess how many of these clinically relevant substitutions lie on the allosteric communication pathway, their spatial positions were identified and mapped onto the 3D structures of TEM-1 and KPC-2. The site of the mutation was plotted as a sphere on its unique Cα position on the structure (Figure 6), which was rendered to represent the allosteric communication pathways shown in Figure 4.

**Figure 6:**
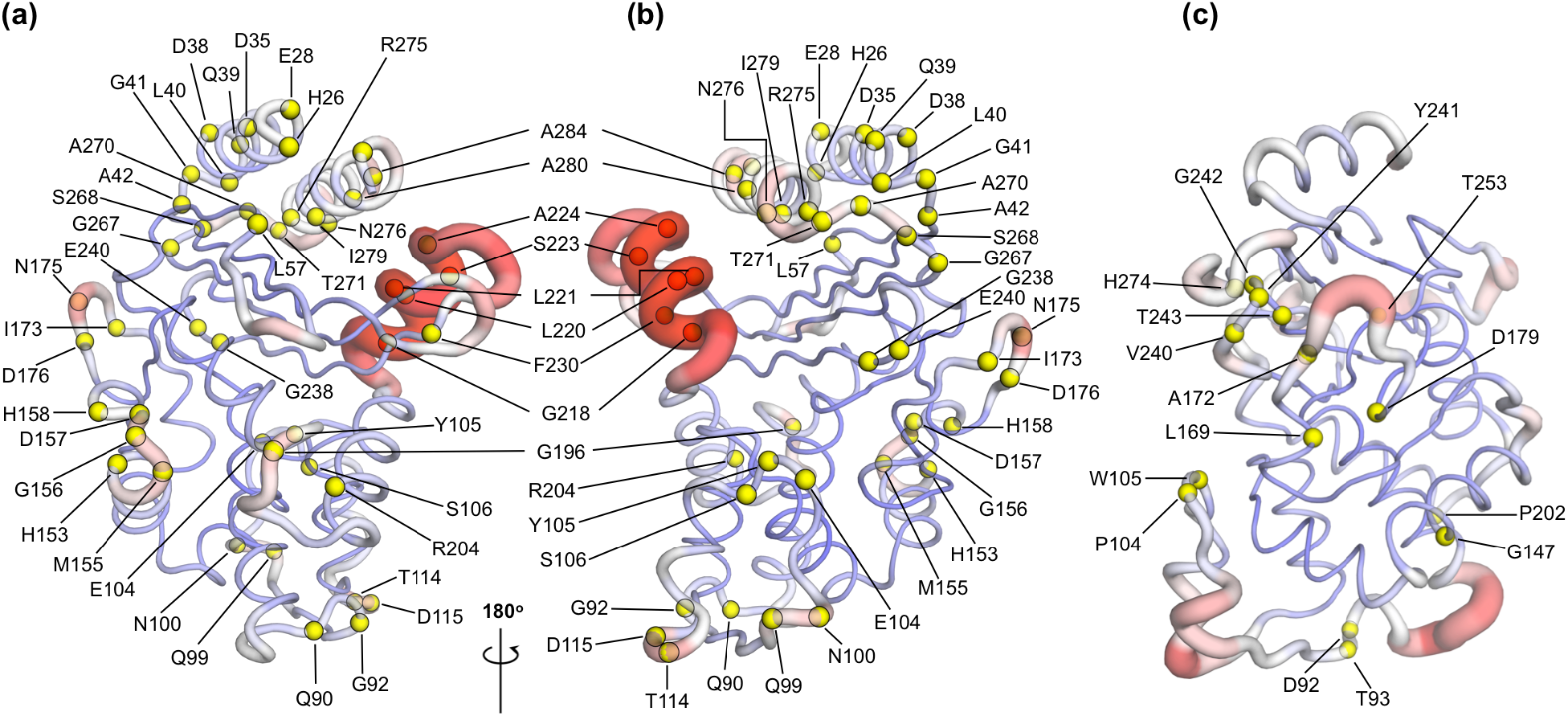
Variant positions in (a, b) TEM-1 and (c) KPC-2 mapped on the averaged Apo_EQ_ structures, also showing allosteric communication pathways (See Figure 4) identified by nonequilibrium simulations. The position of the variant is shown as yellow spheres centered at the corresponding Cα. Only the sites of mutations that lie on the allosteric communication pathways have been annotated. The color scheme and cartoon thickness of the rendered structures represents a snapshot of average Cα deviation between IB_EQ_ and Apo_NE_. Many of these clinically important variant positions lie on the allosteric communication pathway: 45 of the 90 for TEM-1, 15 out of the 25 for KPC-2 single point variants lie on the pathways. This suggests that these variations affect the allosteric behavior of the enzymes.

For TEM-1, 45 of the 90, and for KPC-2 15 out of the 25, amino acid positions known to vary in clinical isolates could be mapped on the allosteric communication pathway. Notably, in TEM-1, residues such as Gly92 preceding b_4_, His153 at the end of a_7_, and Ala224 preceding a_11_; have all been associated with ESBL and/or inhibitor resistant phenotypes identified in the clinic. ^8^. Residues such as M182 and A184, which precede a_9_ and are not on the communication pathway per se; are however surrounded on all sides by loops that are involved in the communication network (Figure S16). For KPC enzymes, where less information is available, characterized variants that have emerged in the clinic differ mostly in activity towards ceftazidime ^58^ and feature substitutions at positions (104, 240, 274) closer to the active site. As more sequences emerge and their phenotypic consequences are described, however, it will then be of interest to establish the properties of KPC variants featuring substitutions at positions (e.g. 92, 93), which lie along the communication pathways described here. We propose that some of these variants differ in allosteric properties, and further, that these differences relate to variances in their clinically relevant spectrum of activity. If our hypothesis is correct, 50% or more of known clinically important variants in these two enzymes may differ in their allosteric behavior, indicating that this is a fundamentally important property. The relationship between sequence (especially substitutions remote from the active site), protein dynamics, spectrum of activity, catalytic turnover and, potentially, allosteric behavior, will be an important future direction in understanding antimicrobial resistance due to β-lactamase enzymes.

## Conclusions

Here, we have identified structural communication between two allosteric binding sites and structural elements, close to the active site, that control enzyme specificity and activity in two distinct, clinically important, class A β-lactamases. The extensive equilibrium MD simulations, with and without ligands, reveal ligand-induced conformational changes, while nonequilibrium MD simulations show that changes at allosteric sites are transmitted to the active site, and identify the structural pathways involved. These nonequilibrium simulations identify the initial stages of the dynamic rearrangement of secondary structural elements and identified the signal propagation routes (with demonstration of its statistical significance). These two complementary approaches together facilitate understanding of how information flows from one part of the protein structure to another.

The equilibrium simulations (of ligand-bound and Apo enzymes) show that the structural effects of ligand binding to allosteric sites are not restricted to the local binding pocket. Class A β-lactamases are rigid enzymes ^31^ that do not undergo large-scale conformational changes; the observed structural rearrangements (caused by ligand removal) are dominated by localized changes in the conformation of loops. Such ligand-induced structural changes are observed in the loops surrounding the active sites including the hinge region, the Ω loop and the α_3_-turn-α_4_ helix, positioned as far as ∼33 Å from the allosteric ligand binding site. In both enzymes, the observed flexible motions lead to an enlargement of the active site, with the potential consequences for the orientation of either mechanistically important regions of the protein or of bound ligand, and, consequently, enzyme activity. It is noteworthy that, while the collective motions associated with the signal propagation were sampled, the PCA does not show the temporal order of events.

The nonequilibrium simulations, using an emerging technique, identify the structural rearrangements arising as a result of a perturbation (ligand removal) and demonstrate communication between the allosteric site and the active site. The ordering of these conformational changes shows the initial steps of communication between secondary structure elements. This structural relay constitutes a pathway that enables effective signal propagation within the enzymes. In TEM-1, the conformational changes initiated at the allosteric site (which is situated between helices α_11_-α_12_) proceed via the β_1_-β_2_ loop to the α_9_-α_10_ loop. From this point, the signal bifurcates towards the Ω loop via the α_7_-α_8_ loop or towards the α_3_-α_4_ pivot via the α_2_-β_4_ loop. In KPC-2, the perturbation caused by ligand unbinding between the α_2_ and α_7_ helices results in conformational changes in loop α_2_-β_4_, leading to β_4_ and onwards to the pivot of the α_3_-turn-α_4_ helix. These conformational changes are relayed to the Ω loop via the α_7_-α_8_ loops. In addition, the signal can also take another route from the α_7_-α_8_ loop towards the β_9_-α_12_ loop, which lies adjacent to the hinge region. It is worth emphasizing that the TEM-1 and KPC-2 systems display a striking resemblance in that the flow of information is towards a common endpoint, despite the two different points of origin. Thus, even though the propagation pathway taken is different, in each case, the signals accumulate to have a structural impact on the conformation of the Ω loop and the α_3_-turn-α_4_ helix. These results demonstrate communication between allosteric ligand binding sites and the active sites of the enzymes, which could be exploited in alternative strategies for inhibitor development.

All class A β-lactamase enzymes share conserved structural architecture.^30,47^ Mutational studies and the location of sites of substitutions in clinical variants suggest the importance to activity of the hinge region, Ω loop and α_3_-turn-α_4_ helix, including the spatial position of the conserved aromatic residue at 105 (or the analogous position in other class A β-lactamases).^8,30,36,66^ Perturbations around these sites, as identified in the simulations here, may constitute a general mechanism by which a conformational signal, transmitted from an allosteric site is relayed via cooperative coupling of loop dynamics to affect catalytic activity. Exploitation of such signaling networks may constitute a novel strategy for the development of new types of inhibitors for these key determinants of bacterial antibiotic resistance.

## Supporting information

movie_S1-TEM1

movie_S2-TEM1

movie_S3-KPC2

## ASSOCIATED CONTENT

### Supporting Information

Structural nomenclature, Dynamical properties, Catalytic cycle, Schematic description of long equilibrium and short nonequilibrium simulations, Time series of Cα RMSD, Core RMSD, Radius of gyration, Solvent accessible surface area, Secondary structure elements, Principal component analysis, PC of TEM-1, PC of KPC-2, Positional Cα RMSF, Ligand dynamics, Cα deviation from subtraction method, TEM-1 DCCM, KPC-2 DCCM, Spatial positions of M182, A184.

## Corresponding Author

Shozeb Haider

