## Supplementary File for "Allosteric communication in Class A β-lactamases occurs via Cooperative Coupling of Loop Dynamics"

**Supporting Information**

**for**

**Publication**

Contents:


| | | |
|---|---|---|
| | Methods | Page S3 |
| Table S1. | Structural nomenclature | Page S8 |
| Table S2. | Dynamical properties | Page S9 |
| Figure S1. | Catalytic cycle of a class A β-lactamase | Page S10 |
| Figure S2. | Schematic description of simulations | Page S11 |
| Figure S3. | Time series of Cα RMSD | Page S12 |
| Figure S4. | Core RMSD | Page S13 |
| Figure S5. | Radius of gyration | Page S14 |
| Figure S6. | Solvent accessible surface area | Page S15 |
| Figure S7. | Secondary structure elements | Page S16 |
| Figure S8. | Principal component analysis | Page S17 |
| Figure S9. | PC of TEM-1 | Page S18 |
| Figure S10. | PC of KPC-2 | Page S19 |
| Figure S11. | Positional Cα RMSF | Page S20 |
| Figure S12. | Ligand dynamics | Page S21 |
| Figure S13. | Cα deviation from subtraction method | Page S22 |
| Figure S14. | TEM-1 DCCM | Page S23 |
| Figure S15. | KPC-2 DCCM | Page S24 |
| Figure S16. | Spatial positions of M182, A184 | Page S25 |
| Figure S17. | Movies: | Page S26 |

**Methods**

To study allosteric modulation of class A β-lactamases, we started by identifying crystal structures of TEM-1 and KPC-2 β-lactamases with allosteric ligands bound. From the ~80 structures present in the Protein Data Bank (PDB), there are only two crystal structures of class A β-lactamases that have a ligand bound in an allosteric pocket. For TEM-1, the 1.45 Å crystal structure in complex with FTA [3-(4-phenylamino-phenylamino)-2-(1h-tetrazol5-yl)-acrylonitrile] was chosen as the starting structure (PDB id: 1PZP) for this work.[1] In this structure, the inhibitor binds between helices $\alpha_{11}$ and $\alpha_{12}$ (Figure 1a), in a site ~16 Å away from the active site Ser70. Two unstructured residues from the C-terminal end (His289, Trp290) were removed from the crystal structure. For KPC-2, the 1.35 Å crystal structure in complex with a coumarin phosphonate analogue, GTV [(5,7-dimethyl-2-oxo-2h-1-benzopyran-4-yl)methyl]phosphonic acid], was chosen as the starting structure (PDB id: 6D18).[2] GTV binds in three sites on KPC-2 (Figure 1b): the first is in the active site (orthosteric ligand); the second site is adjacent to helix $\alpha_6$ (allosteric ligand 1); and the third (allosteric ligand 2) is on the distal end of the enzyme, ~16 Å from the active site Ser70 in between helices $\alpha_2$ and $\alpha_7$. The orthosteric and allosteric ligand1 (Figure 1b) were discarded because of their direct proximity to the active site and replaced by water. Three unstructured residues from the N-terminal end (His23, Met24, Leu25) and seven from the C-terminal end (Leu288-Gly294) were removed from the starting structure to avoid any simulation artifacts arising as a result of terminal fraying during simulations.

The ionization states of the amino acid side chains were determined at pH 7.0, using the *ProteinPrepare* functionality as implemented in the High-Throughput

Molecular Dynamics (HTMD) framework.[3,4] Charges were assigned on the basis of their local environment, via optimization of the hydrogen-bonding network of the protonated structure.[3]

Parameters for the ligands were generated using the Antechamber tool.[5] The geometry was optimized at the B3LYP/6-31G(d) level and RESP charges were fitted using electrostatic potential obtained at the HF/6-31G(d) level.[6] The necessary nonbonded parameters for the dynamics of the ligands were adopted from GAFF2.[7] All complexes were set up using tleap, as implemented in the Amber MD package.[8] The Amber ff14SB forcefield [9] was used for the protein. In total, four complexes were set up, including an allosteric inhibitor-bound (IB) and an Apo (no ligand) system for both TEM-1 and KPC-2 β-lactamases. The Apo system was generated by removing the inhibitor from the allosteric binding site. In all simulated complexes, there is no ligand bound to the orthosteric site. Each complex was solvated using TIP3P water in a cubic box, whose edge was set to at least 10 Å from the closest solute atom.[10] The systems were neutralized using $K^+$ and $Cl^-$ counter ions. The simulation protocol was identical for each system. The systems was minimized and relaxed under NPT conditions for 5 ns at 1 atm. The temperature was increased to 300 K using a timestep of 4 fs, rigid bonds and a cutoff of 9 Å and particle mesh Ewald summations switched on for long-range electrostatics.[11] During the equilibration step, the protein's backbone and the ligand atoms were restrained by a spring constant set at 1 kcal $mol^{-1}$ $Å^{-2}$, while the ions and solvent were free to move. The production simulations were run in the NVT ensemble using a Langevin thermostat with a damping constant of 0.1 ps and hydrogen mass repartitioning scheme to achieve a time step of 4 fs. [12] The final production step was run without any restraints. All simulations were run using the

ACEMD molecular dynamics engine as implemented in the HTMD framework.[4] Visualization of the simulations was done using the VMD package.[13]

*Equilibrium simulations:* In order to sufficiently sample the conformational space, 20 replicate simulations of 250 ns each were performed for each system. This resulted in a total sampling time of 5 μs for each system. The initial velocities of the atoms of each replica were randomized. We address this set of runs in this study as *equilibrium simulations* ($Apo_{EQ}$/$IB_{EQ}$).

*Nonequilibrium simulations:* To investigate rapid conformational changes and study signal propagation within the proteins, we carried out 800 short nonequilibrium MD simulations for each system. Such nonequilibrium simulations have been applied successfully to study inter-domain communication in receptors and other systems like ABC transporters.[14–18] We used the Kubo-Onsager approach [19–21] to extract the conformational response of the proteins to ligand removal. In this approach, the response of a system to a perturbation is computed by calculating the difference in the evolution of the simulations with and without the perturbation. Subtracting the perturbed and unperturbed pairs of simulations at a given time, and averaging the results over multiple replicates, allows not only for the identification of the events associated with signal propagation but also determines the statistical significance of the observations. When the two sets of simulations (with and without a perturbation) are correlated, the subtraction technique permits the cancellation of noise arising from random intrinsic fluctuations of the system thus allowing the identification of the response to the perturbation in a statistically significant way.[21] In our systems, the perturbation was generated by (instantaneously) removing the ligand from the allosteric pocket. It is important to emphasize that the annihilation of the ligand in this way does not represent the physical process of unbinding. The objective is to rapidly elicit response and force signal propagation within the protein, as the conformation adjusts to the removal of the ligand. Such a response allows for the identification of the initial signals that are sent out as conformational changes associated with the signal propagating from the allosteric binding pocket. The structural rearrangements in the communication pathways revealed by nonequilibrium simulations are likely to be involved in response to the physical process of binding and unbinding of ligands in the allosteric pockets.

A graphical representation of the procedure that was followed to setup the nonequilibrium simulations is given in Figure S2. The starting conformation for the short nonequilibrium simulations ($Apo_{NE}$) was extracted from the equilibrated part of the 250 ns equilibrium simulations (50-250 ns). Specifically, conformations were taken every 5 ns, the ligand was removed from the allosteric pocket and the resulting $Apo_{NE}$ system was run for another 5 ns (Figure S2). 40 short, nonequilibrium simulations were run for each replicate. In total 800 simulations were run for each system. The simulation conditions of the nonequilibrium simulations were identical to those in the equilibrium simulations.

For each pair of unperturbed $IB_{EQ}$ and perturbed $Apo_{NE}$ simulations, the difference in positions for each C$\alpha$ was determined at equivalent points in time, namely at 0, 0.05, 0.5, 1, 3 and 5 ns. Calculating the differences in the positions of C$\alpha$ identifies conformational rearrangements, while reducing the noise coming from side chain fluctuations. The C$\alpha$ deviation values at each time point were were averaged over all 800 simulations. The statistical significance of the structural changes is demonstrated by the low standard error of the averages.

The analysis was carried out using Gromacs tools [22], MDLovofit [23] and in-house scripts.[18] Principal Component Analysis was carried out to explore the conformational space that the trajectories sampled and identify the motions in the protein with the highest amplitude using PyPcazip.[24,25] The $Apo_{EQ}$ and $IB_{EQ}$ trajectories were combined so that all share the same subspace and comparisons can be made. The first two principal components (PC1 and PC2) were used to assess the equilibration of the simulation, as the first few principal components represent the slower intrinsic large-scale motions of the protein.[26] All systems were considered equilibrated after 50 ns. The dynamic cross-correlations for Cα-Cα were calculated using cpptraj analysis program.[27] The results were plotted using in-house scripts and visualized using MATLAB (www.mathworks.com).

An independent-samples Student's t-test was used to compare the $Apo_{EQ}$ and $IB_{EQ}$ RMSFs and to assess the significance of the differences observed.[18,28] The sample size used for the t-test was the 20 RMSF profiles of the $Apo_{EQ}$ and $IB_{EQ}$ independent simulations. The assumption used for the t-test was that the samples from the two states were independent, the dependent variable was normally distributed and the variances of the dependent variable were equal.

The figures were made using PyMol, VMD [13], ChimeraX [29] and Molsoft ICM-Pro package (www.molsoft.com). The manuscript was written in the Manuscripts app (www.manuscripts.io).

**Table S1: Structural nomenclature**. The loops are named based on the secondary structure it connects. For example, loop $\alpha_1$-$\beta_1$ connects $\alpha_1$ helix and $\beta_1$ sheet. It must be noted that the boundaries of the secondary structure are approximate and may vary by ±2 residues based on the visualization software used. This work employed ChimeraX to define secondary structure. The mutations listed are those that fall on the communication pathway.

| Secondary Structure | Residues | | | |
|---|---|---|---|---|
| | **TEM-1 (PDB id 1PZP)** | TEM-1 Mutations | **KPC-2 (PDB id 6D18)** | KPC-2 Mutations |
| $\alpha_1$ | 26-40 | H26, E28, D35, D38, Q39, L40, | 26-40 | |
| $\alpha_1$-$\beta_1$ | 41-42 | G41, A42 | 41-42 | |
| $\beta_1$ | 43-50 | | 43-50 | |
| $\beta_1$-$\beta_2$ | 51-55 | | 51-55 | |
| $\beta_2$ | 56-60 | L57 | 56-60 | |
| $\beta_2$-$\beta_3$ | 61-65 | | 61-65 | |
| $\beta_3$ | 66-67 | | 66-67 | |
| $\beta_3$-$\alpha_2$ | 68-71 | | 68-70 | |
| $\alpha_2$ | 72-85 | | 72-87 | |
| $\alpha_2$-$\beta_4$ | 86-93 | Q90, G92 | 88-93 | D92, T93 |
| $\beta_4$ | 94-95 | | 94-95 | |
| $\beta_4$-$\alpha_3$ | 96-97 | | 96-97 | |
| $\alpha_3$ | 98-101 | Q99, N100, | 98-102 | |
| $\alpha_3$-turn-$\alpha_4$ | 102-107 | E104, Y105, S106 | 103-107 | P104, W105 |
| $\alpha_4$ | 108-111 | | 108-113 | |
| $\alpha_4$-$\beta_5$ | 112-116 | T114, D115 | 114-116 | |
| $\beta_5$ | 117-118 | | 117-118 | |
| $\alpha_5$ | 119-129 | | 119-129 | |
| $\alpha_5$-$\alpha_6$ | 130-131 | | 130 | |
| $\alpha_6$ | 132-142 | | 131-142 | |
| $\alpha_6$-$\alpha_7$ | 143-144 | | 143-144 | |
| $\alpha_7$ | 145-154 | H153 | 145-155 | G147 |
| $\alpha_7$-$\alpha_8$ | 155-166 | M155, G156, D157, H158 | 156-166 | |
| $\alpha_8$ | 167-171 | | 167-171 | L169 |
| Ω | 172-179 | I173, N175, D176 | 172-179 | A172, D179 |
| $\beta_6$ | 180-181 | | 180-181 | |
| $\beta_6$-$\alpha_9$ | 182 | | 182 | |
| $\alpha_9$ | 183-195 | | 183-195 | |
| $\alpha_9$-$\alpha_{10}$ | 196-200 | G196 | 196-199 | |
| $\alpha_{10}$ | 201-212 | R204, | 200-213 | P202 |
| Hinge | 213-218 | G218 | 214-218 | |
| $\alpha_{11}$ | 219-224 | L220, L221, S223, A224 | 219-225 | |
| $\alpha_{11}$-$\beta_7$ | 225-229 | | 226-229 | |
| $\beta_7$ | 230-237 | F230 | 230-237 | |
| $\beta_7$-$\beta_8$ | 238-243 | G238, E240 | 238-243 | V240, Y241, G242, T243 |
| $\beta_8$ | 244-251 | | 244-251 | |
| $\beta_8$-$\beta_9$ | 252-258 | | 252-258 | T253 |
| $\beta_9$ | 259-266 | | 259-266 | |
| $\beta_9$-$\alpha_{12}$ | 267-272 | G267, S268, A270, T271, | 267-273 | |
| $\alpha_{12}$ | 273-288 | R275, N276, I279, A280, A284 | 274-287 | H274 |

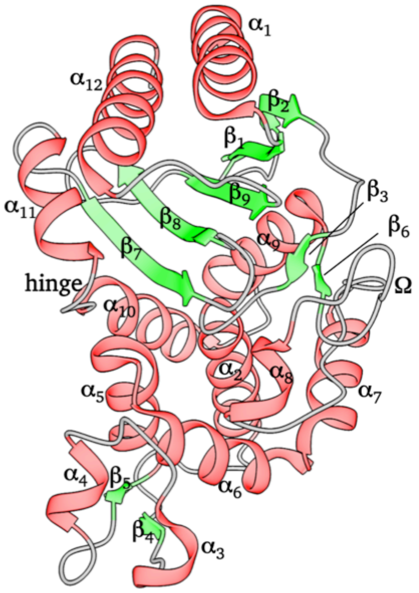

**Table S2:** Dynamical properties (RMSD, Rg and SASA) used to assess structural stability of the systems over the course of the equilibrium simulation. The values represent the averages calculated from the equilibrated part (50-250 ns) of all 20 replicate simulations of each system.

| Property | System | | Value |
|---|---|---|---|
| RMSD | TEM-1 | $Apo_{EQ}$ | 0.12 nm |
| | | $IB_{EQ}$ | 0.11 nm |
| | KPC-2 | $Apo_{EQ}$ | 0.11 nm |
| | | $IB_{EQ}$ | 0.10 nm |
| Rg | TEM-1 | $Apo_{EQ}$ | 1.78 nm |
| | | $IB_{EQ}$ | 1.79 nm |
| | KPC-2 | $Apo_{EQ}$ | 1.78 nm |
| | | $IB_{EQ}$ | 1.78 nm |
| SASA | TEM-1 | $Apo_{EQ}$ | 118 $nm^2$ |
| | | $IB_{EQ}$ | 121 $nm^2$ |
| | KPC-2 | $Apo_{EQ}$ | 114 $nm^2$ |
| | | $IB_{EQ}$ | 115 $nm^2$ |

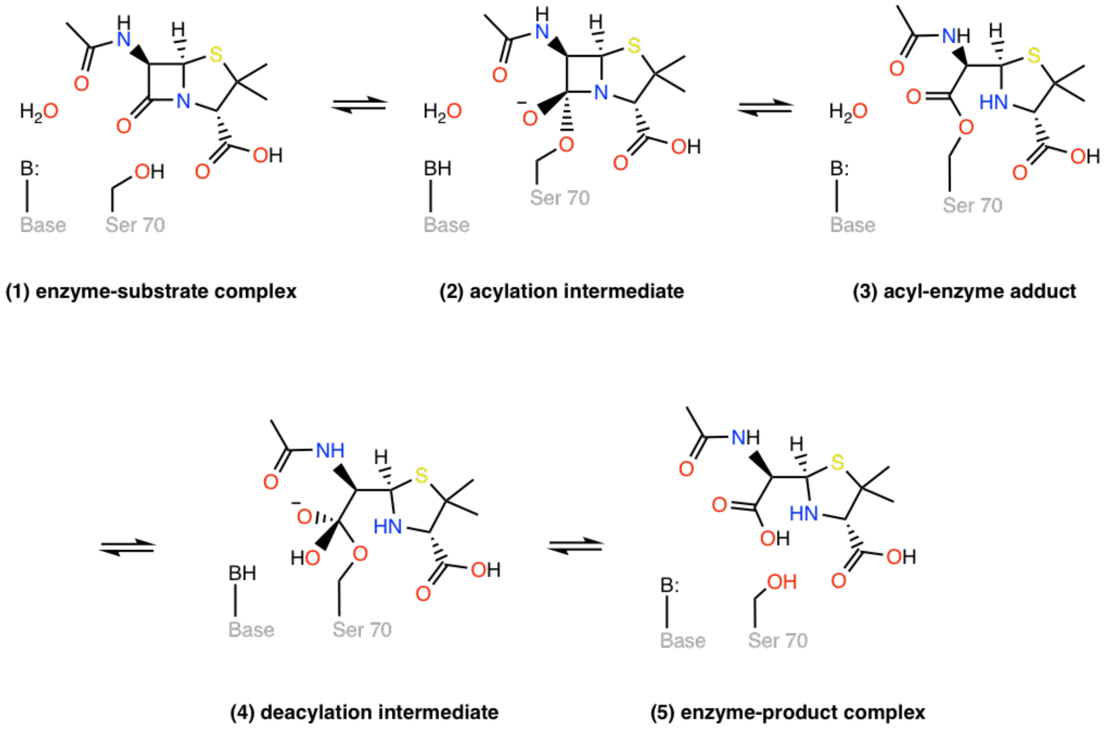


**Figure S1.** Catalytic cycle of a class A β-lactamase illustrated on the core structure of penicillins. Class A β-lactamases use an active site serine nucleophile to cleave the β-lactam bond of the substrate in a two-step acylation-deacylation reaction cycle that leads to overall hydrolysis. (1) The acylation reaction initiates with the reversible binding of the antibiotic in the active site and the formation of the enzyme-substrate complex. In the next step, a general base-catalyzed nucleophilic attack on the β-lactam carbonyl by the serine hydroxyl takes place through a tetrahedral intermediate (2) to form a transient acyl-enzyme adduct (3). In the deacylation step, the acyl-enzyme adduct (3) undergoes a general base-catalyzed attack by a hydrolytic water molecule to form a second tetrahedral intermediate (4), which then forms a postcovalent product complex (5), from which the hydrolyzed product is released.

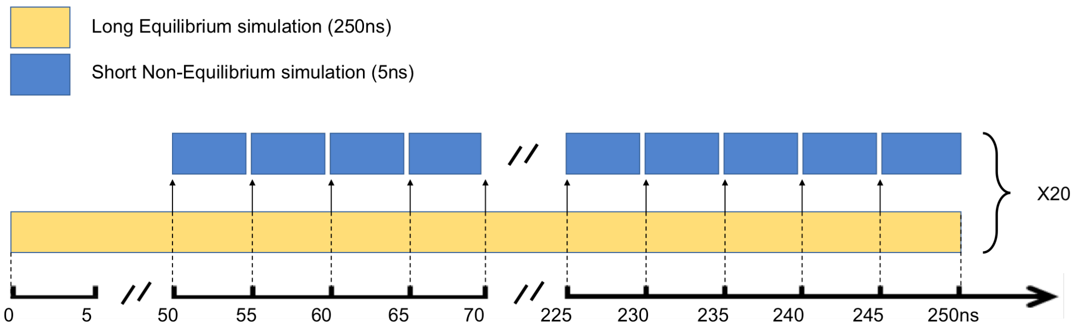


**Figure S2.** Schematic description of the long equilibrium (EQ) and short nonequilibrium (NE) simulations. 20 replicates of $IB_{EQ}$ simulations were run starting from the minimized crystal structure. From the equilibrated part of each replica (50 ns onwards), the final conformation of the protein-ligand complex was extracted at every 5 ns, the perturbation (removal of the ligand) was introduced, and a short $Apo_{NE}$ simulations was run for 5 ns. In total, 800 $Apo_{NE}$ simulations were performed for each system. In addition to these simulations, 20 replicas of the $Apo_{EQ}$ were also simulated.

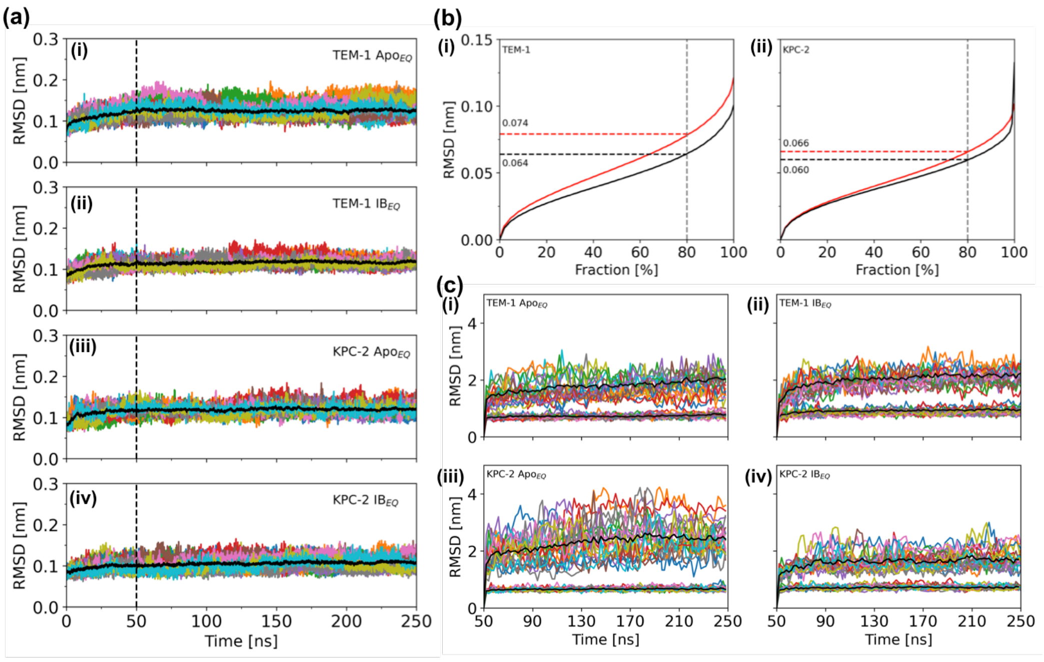


**Figure S3.** (a) Time series of the Cα Root Mean Squared Deviation (RMSD) of (ai) TEM-1 $Apo_{EQ}$, (aii) TEM-1 $IB_{EQ}$, (aiii) KPC-2 $Apo_{EQ}$ and (aiv) KPC-2 $IB_{EQ}$ systems, measured over the course of the 250 ns of each replica. The black line represents the average of the 20 replicates. (b) RMSD calculated as a function of the fraction of the total Cα atoms considered for structural alignment in (i) TEM-1 and (ii) KPC-2. The plots indicate that 80% of conformations in TEM-1 that can be aligned to below 0.064 nm ($Apo_{EQ}$; black) and 0.074 nm ($IB_{EQ}$; red). Similarly, in KPC-2, 80% of the conformations could be superimposed to below 0.060 nm ($Apo_{EQ}$; black) and 0.066 nm ($IB_{EQ}$; red). This constitutes the core of the enzyme. (c) Fractional Cα RMSD calculated after identification of the core in (ci) TEM-1 $Apo_{EQ}$ (cii) TEM-1 $IB_{EQ}$ (ciii) KPC-2 $Apo_{EQ}$ (civ) KPC-2 $IB_{EQ}$. The bottom black line denotes the stable core and consists of 80% Cα atoms. The top black line is the average of the remainder 20% Cα atoms calculated from all 20 replicates. These constitute the non-core Cα atoms that display deviation in all simulations.

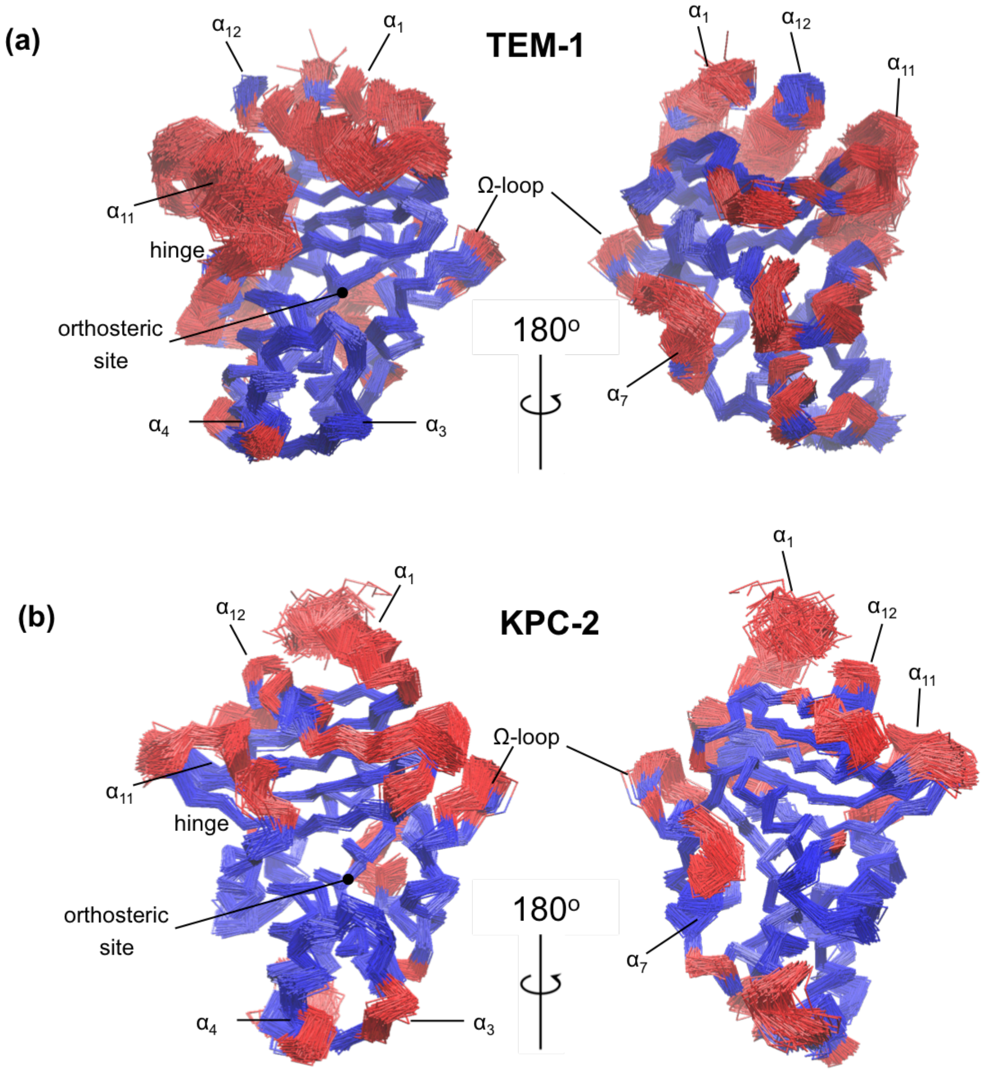


**Figure S4.** Core Cα RMSD superimposition from (a) TEM-1 and (b) KPC-2 $IB_{EQ}$ simulations. The structural alignment was calculated from the equilibrated section of all $IB_{EQ}$ trajectories and rendered to illustrate 100 uniformly separated frames. The least mobile Cα atoms are colored blue and the most mobile atoms (red) provide the structural basis for the differential RMSDs.

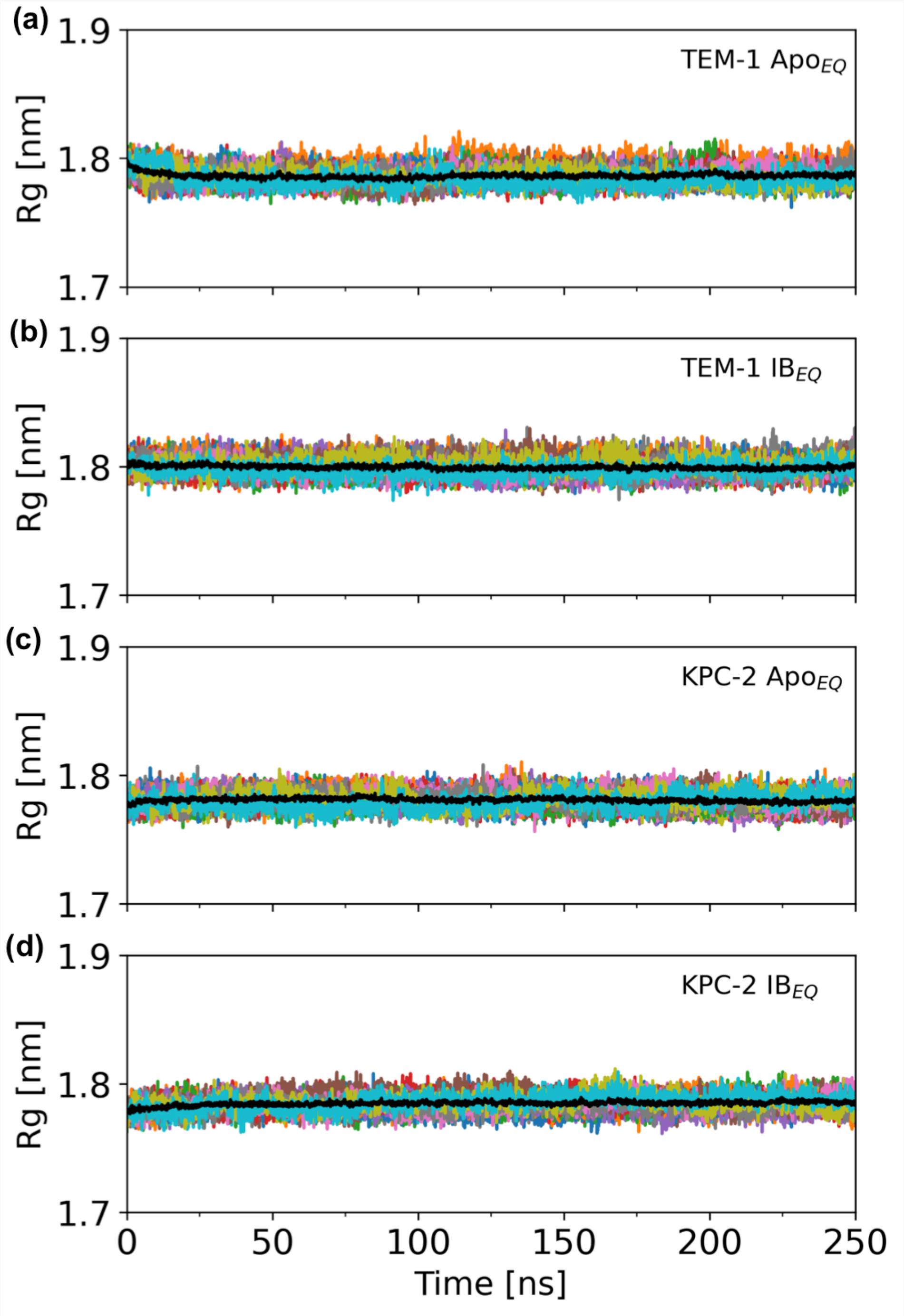


**Figure S5.** Time evolution of the radius of gyration (Rg) of the complete (a) TEM-1 Apo$_{EQ}$, (b) TEM-1 IB$_{EQ}$, (c) KPC-2 Apo$_{EQ}$ and (d) KPC-2 IB$_{EQ}$ enzymes, measured over the course of the 250 ns of each replicate. The black line represents the average of 20 replicates.

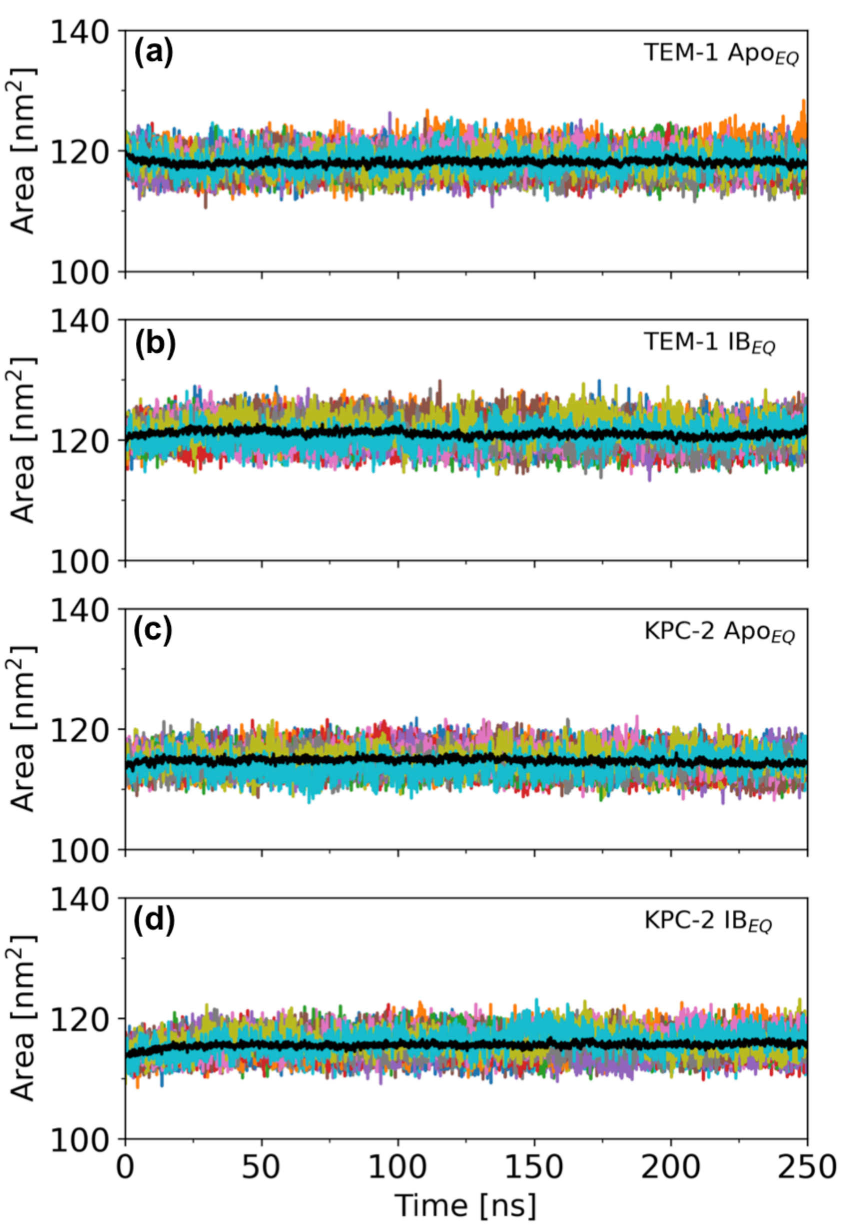


**Figure S6.** Solvent accessible surface area (SASA) was calculated to assess structural distortion in the complete (a) TEM-1 Apo$_{EQ}$, (b) TEM-1 IB$_{EQ}$, (c) KPC-2 Apo$_{EQ}$ and (d) KPC-2 IB$_{EQ}$ enzymes, measured over the course of the 250 ns of each replicate. The black line represents the average of 20 replicates.

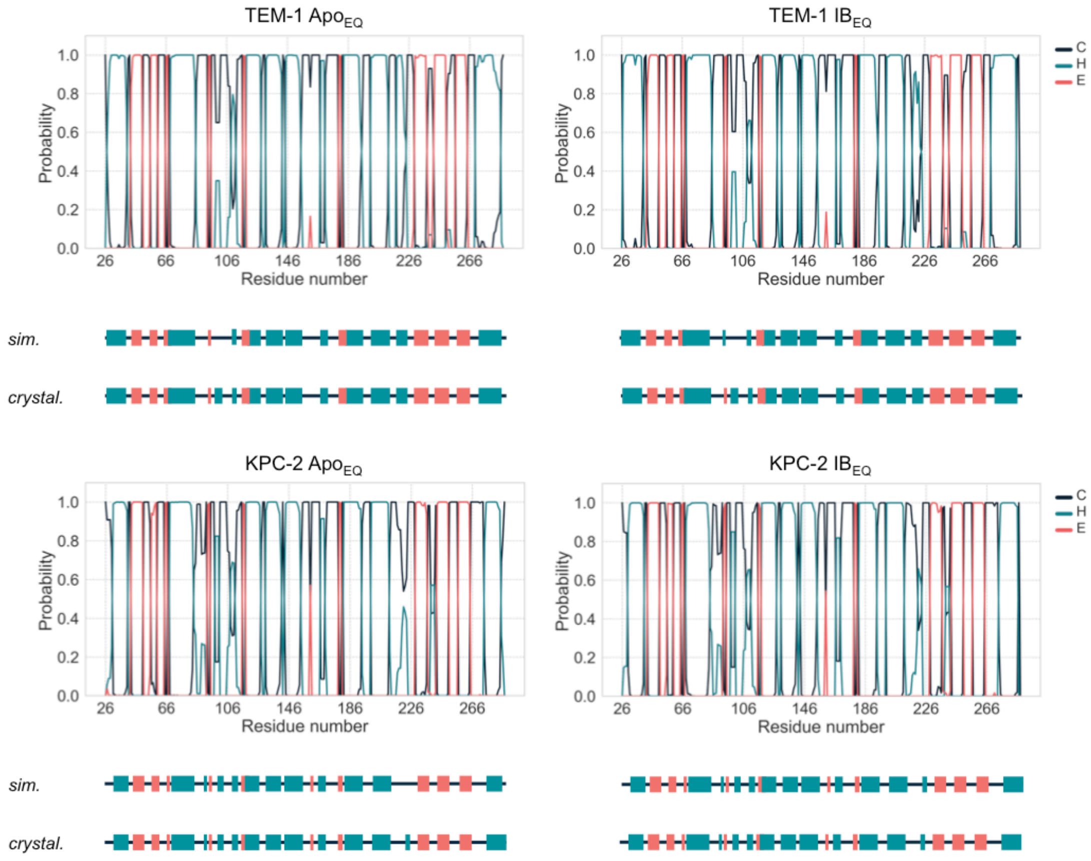


**Figure S7.** Probability to find each residue in a coil ('C'), helix ('H'), or strand ('E'). The secondary structure element was assigned to each residue in each frame of each of the 20 replicas per system using the DSSP algorithm as implemented in MDTraj python library. The secondary structure of TEM-1 and KPC-2 as assigned from the crystal structure (*cryst.*), as well as the most probable assignment according to the simulations (*sim.*) is depicted as a cartoon below each plot.

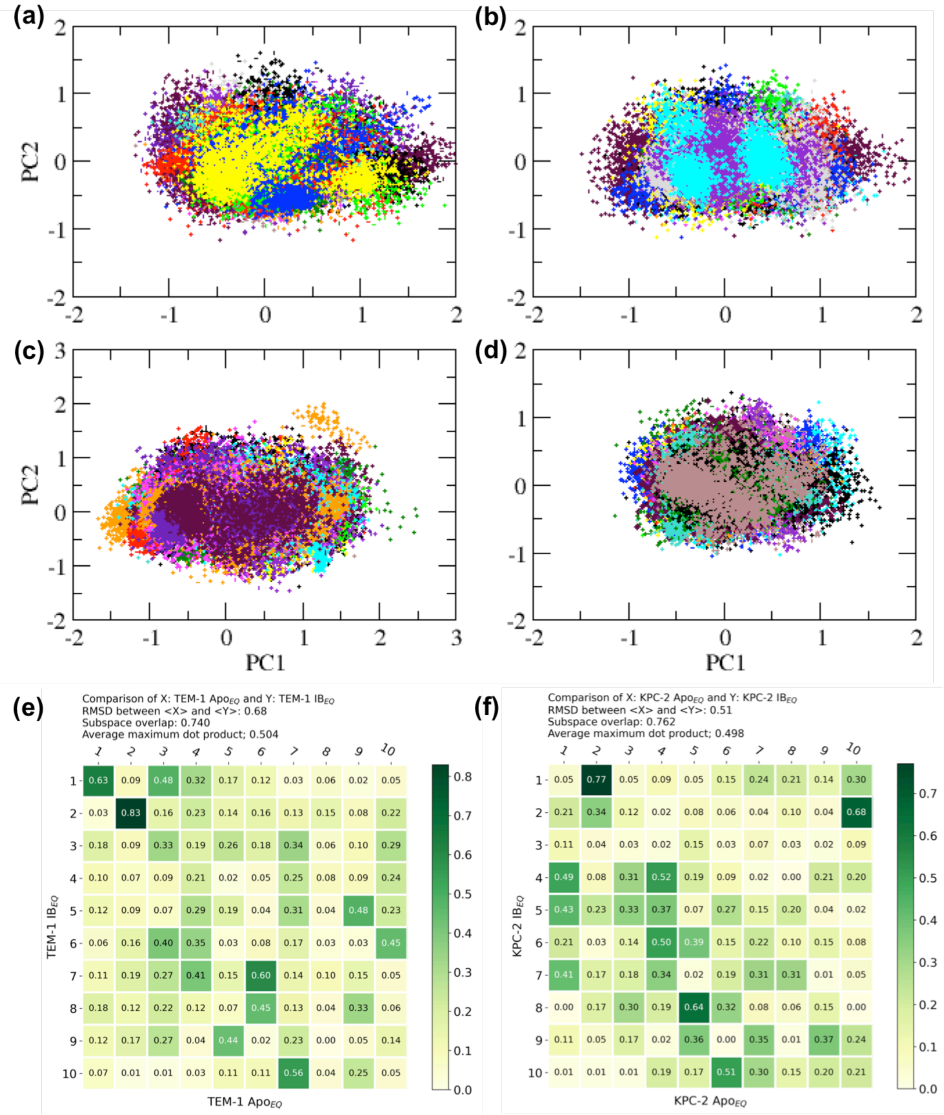


**Figure S8.** Projection of the conformational space sampled in each of the 20 replicates simulations of (a) TEM-1 Apo$_{EQ}$, (b) TEM-1 IB$_{EQ}$, (c) KPC-2 Apo$_{EQ}$ and (d) KPC-2 IB$_{EQ}$ on the first two Principal Components (PC). The subspace that each simulation samples has been highlighted with a different color. A quantitative comparison of the collective modes from (e) TEM-1 Apo$_{EQ}$ and IB$_{EQ}$ and (f) KPC-2 Apo$_{EQ}$ and IB$_{EQ}$ states, is represented as a dot product matrix between the eigenvectors identified from apo and IB states. This dot product highlights the subspace overlap. All eigenvectors were used to calculate the subspace overlap between the Apo$_{EQ}$ and IB$_{EQ}$ states.

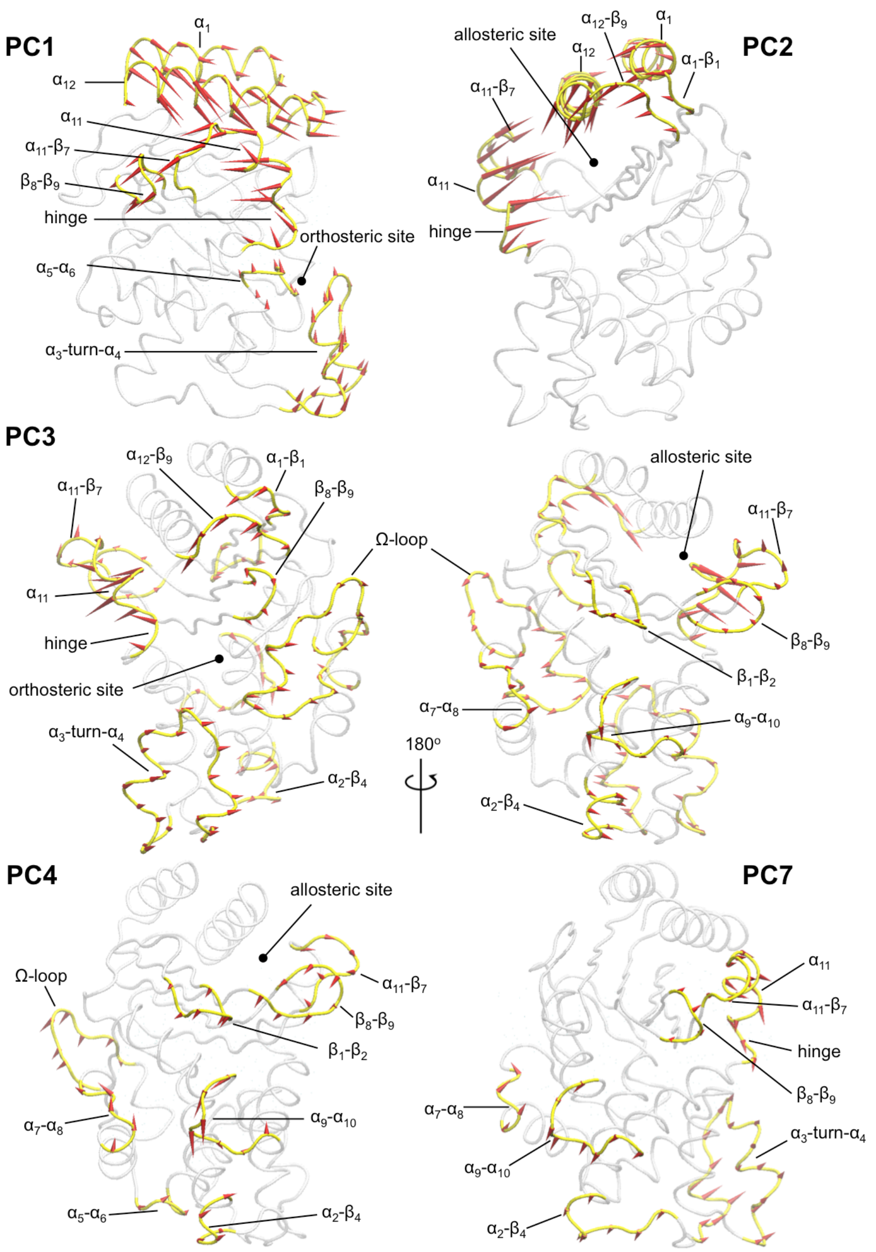


**Figure S9.** Concerted motions observed in TEM-1 equilibrium simulations. For the PCA of equilibrium simulations, all 20 replicates from $\text{Apo}_{\text{EQ}}$ and $\text{IB}_{\text{EQ}}$ were combined before analysis. Motions associated with the enlargement of the active site were observed in PC3. The individual motions linked with signal propagation were sampled in multiple PCs; however, PCA loses the temporal order of events. The red arrows represent the direction and magnitude of the motion.

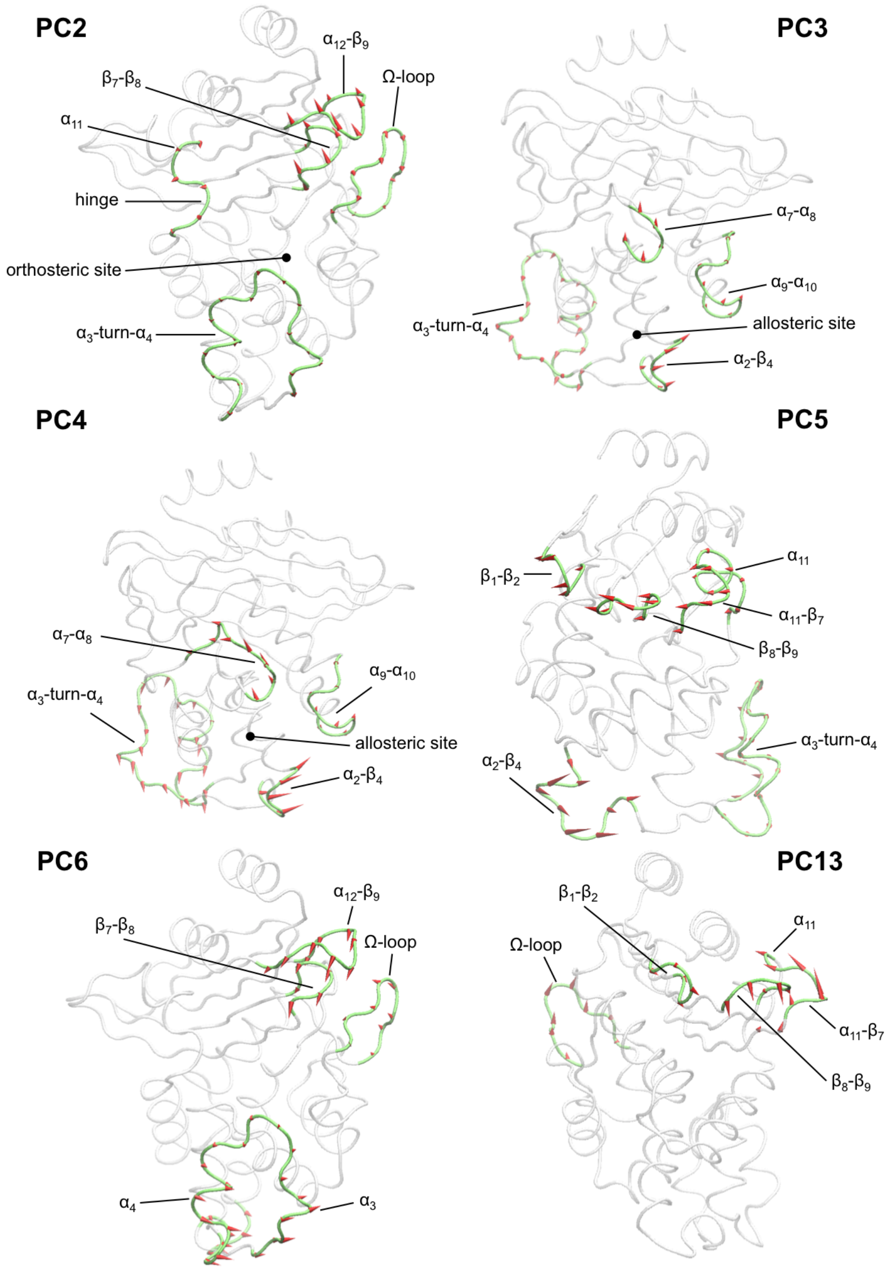


**Figure S10.** Concerted motions observed in KPC-2 equilibrium simulations. For the PCA of equilibrium simulations, all 20 replicates from $Apo_{EQ}$ and $IB_{EQ}$ were combined before analysis. Motions associated with the enlargement of the active site were observed in PC2. The individual motions linked with signal propagation were sampled in multiple PCs; however, PCA loses the temporal order of events. The red arrows represent the direction and magnitude of the motion.

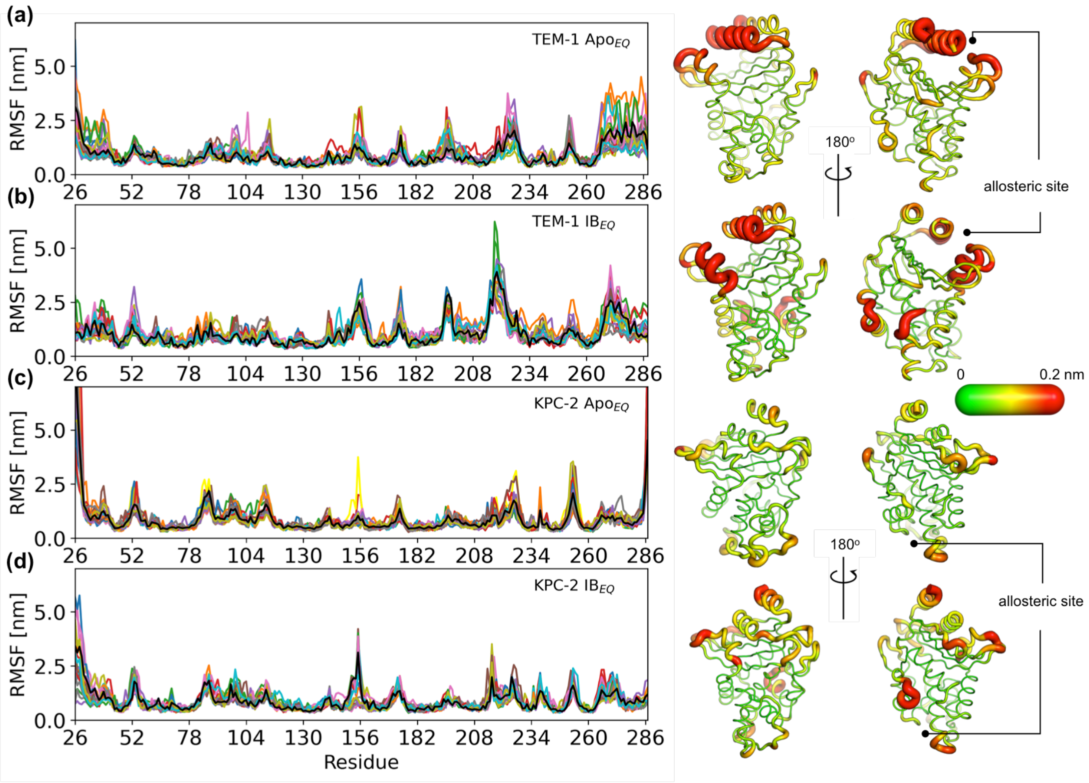


**Figure S11.** Positional Cα Root Mean Square Fluctuation (RMSF) of (a) TEM-1 $Apo_{EQ}$, (b) TEM-1 $IB_{EQ}$, (c) KPC-2 $Apo_{EQ}$ and (d) KPC-2 $IB_{EQ}$ systems. The black line represents the average RMSF calculated from the last 200 ns of all 20 replicate simulations. This average value is mapped on the structure to highlight regions of high flexibility.

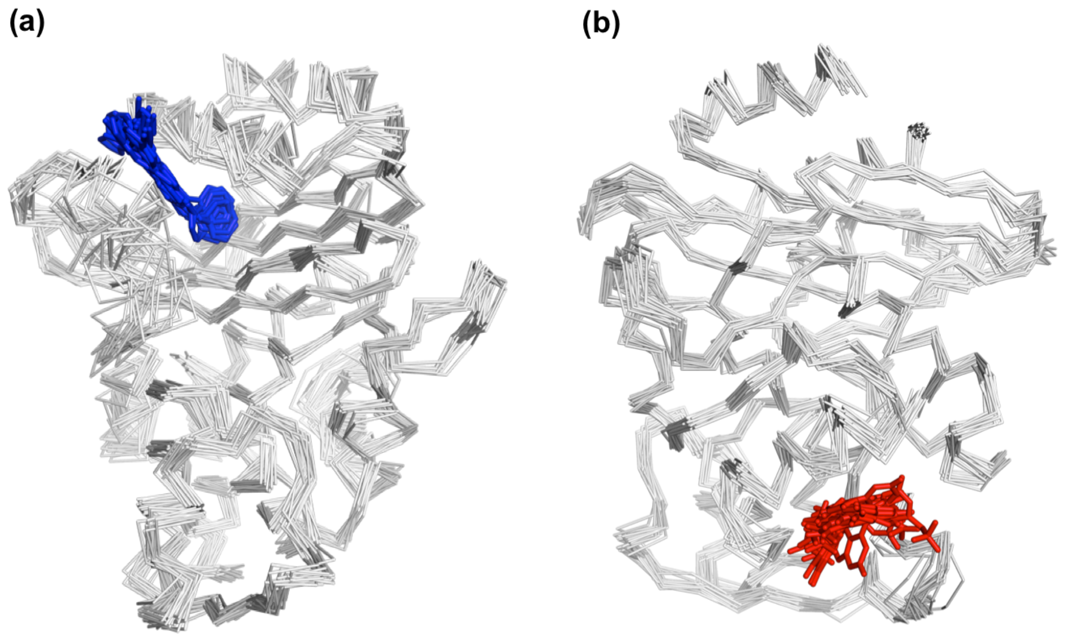


**Figure S12.** Snapshot of the last frame from (a) TEM-1 $IB_{EQ}$ and (b) KPC-2 $IB_{EQ}$ replicate simulations, highlighting the spatial position of the ligands in the allosteric binding sites. FTA is represented as blue sticks and GTV is colored in red.

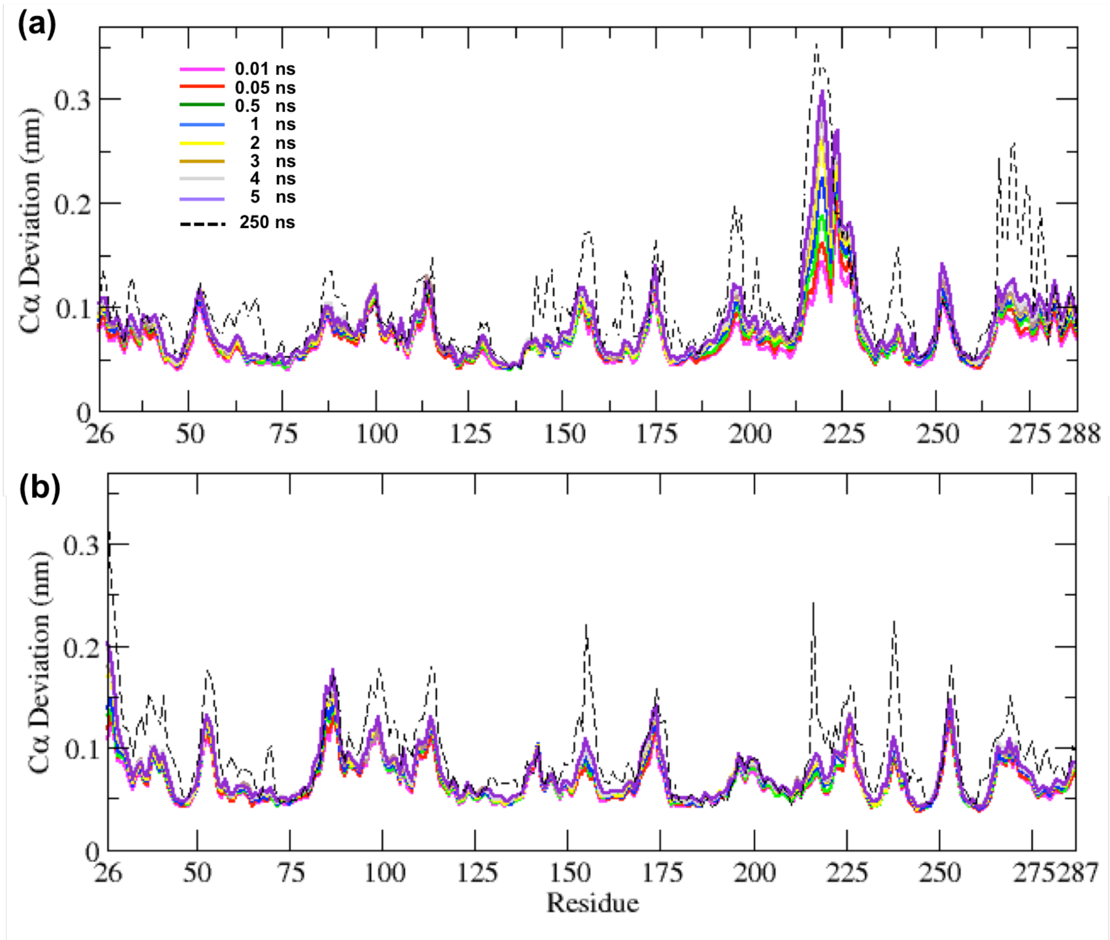


**Figure S13.** Average Cα deviation between the $IB_{EQ}$ and $Apo_{NE}$ calculated using the subtraction method for (a) TEM-1 and (b) KPC-2. Average from all 800 simulations at various time points are illustrated. The average Cα deviation between the $IB_{EQ}$ and $Apo_{NE}$ simulations is plotted as a dotted line for comparison.

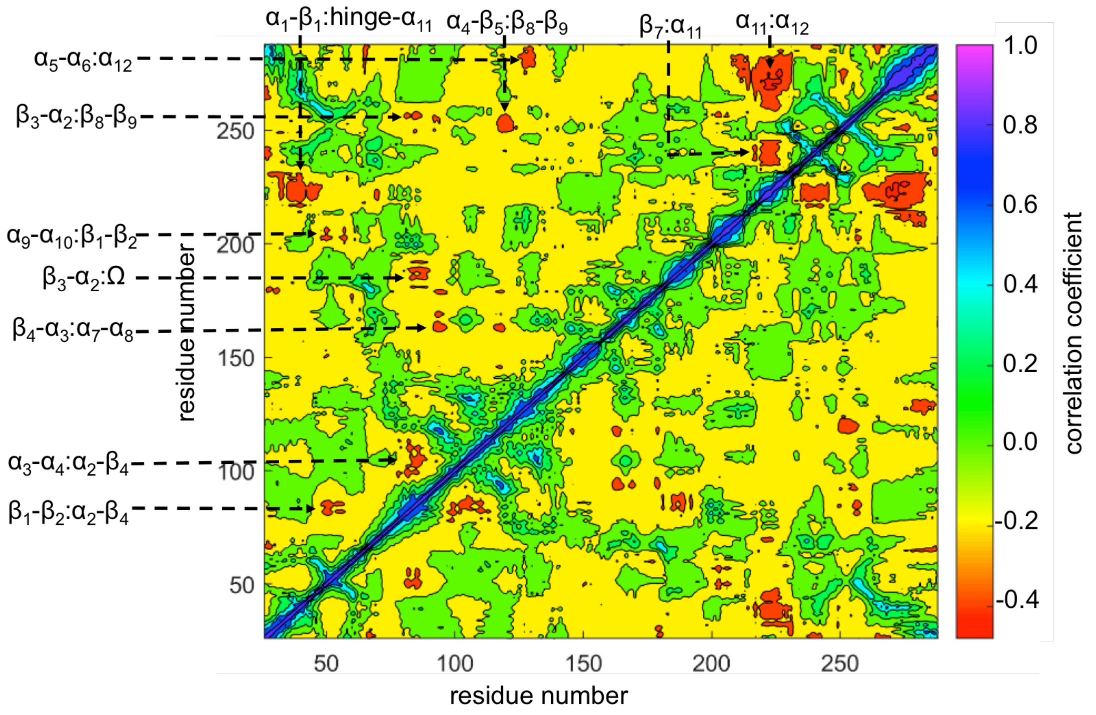


**Figure S14.** TEM-1 averaged DCCM computed from all nonequilibrium trajectories. The regions showing significant correlations are identified: $\beta_1$-$\beta_2$:$\alpha_2$-$\beta_4$, $\alpha_3$-$\alpha_4$:$\alpha_2$-$\beta_4$, $\beta_4$-$\alpha_3$:$\alpha_7$-$\alpha_8$, $\beta_3$-$\alpha_2$:$\Omega$, $\alpha_9$-$\alpha_{10}$:$\beta_1$-$\beta_2$, $\beta_3$-$\alpha_2$:$\beta_8$-$\beta_9$, $\alpha_5$-$\alpha_6$:$\alpha_{12}$, $\alpha_1$-$\beta_1$:hinge-$\alpha_{11}$, $\alpha_4$-$\beta_5$:$\beta_8$-$\beta_9$, $\beta_7$:$\alpha_{11}$ and $\alpha_{11}$:$\alpha_{12}$.

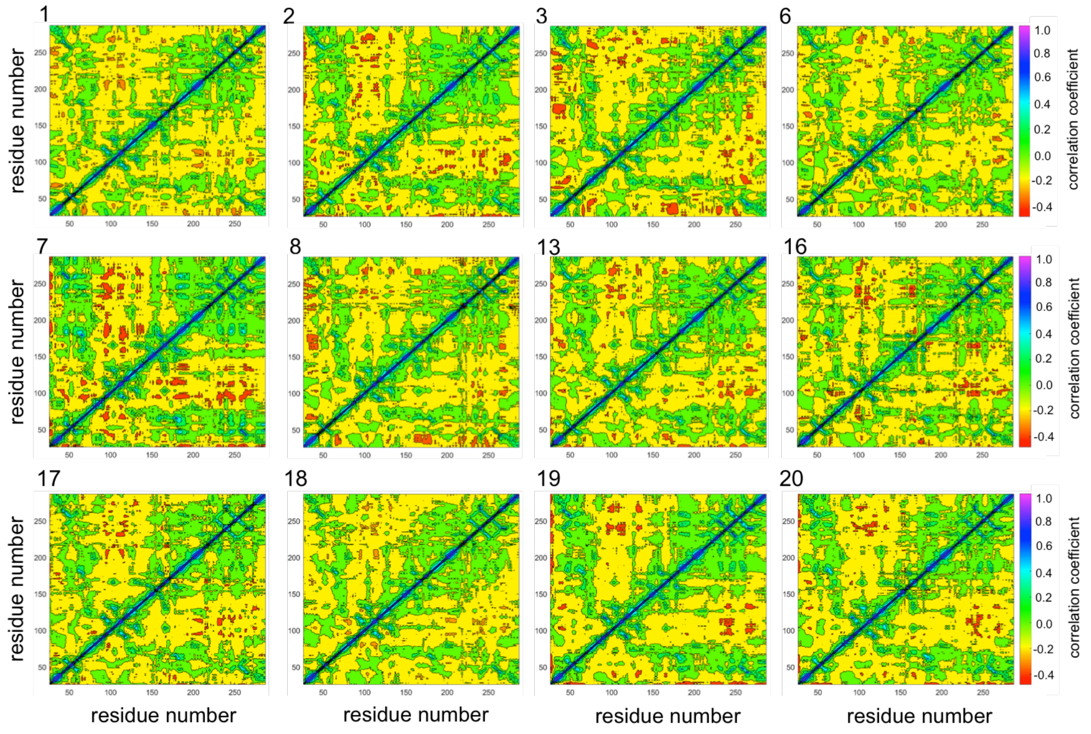


**Figure S15.** Selected DCCMs computed for individual 5 ns nonequilibrium MD trajectories of KPC-2. These individual trajectories show different behavior from the averaged results (see Figure 5), as these trajectories show several regions of significant correlations similar to the TEM-1 nonequilibrium DCCM.

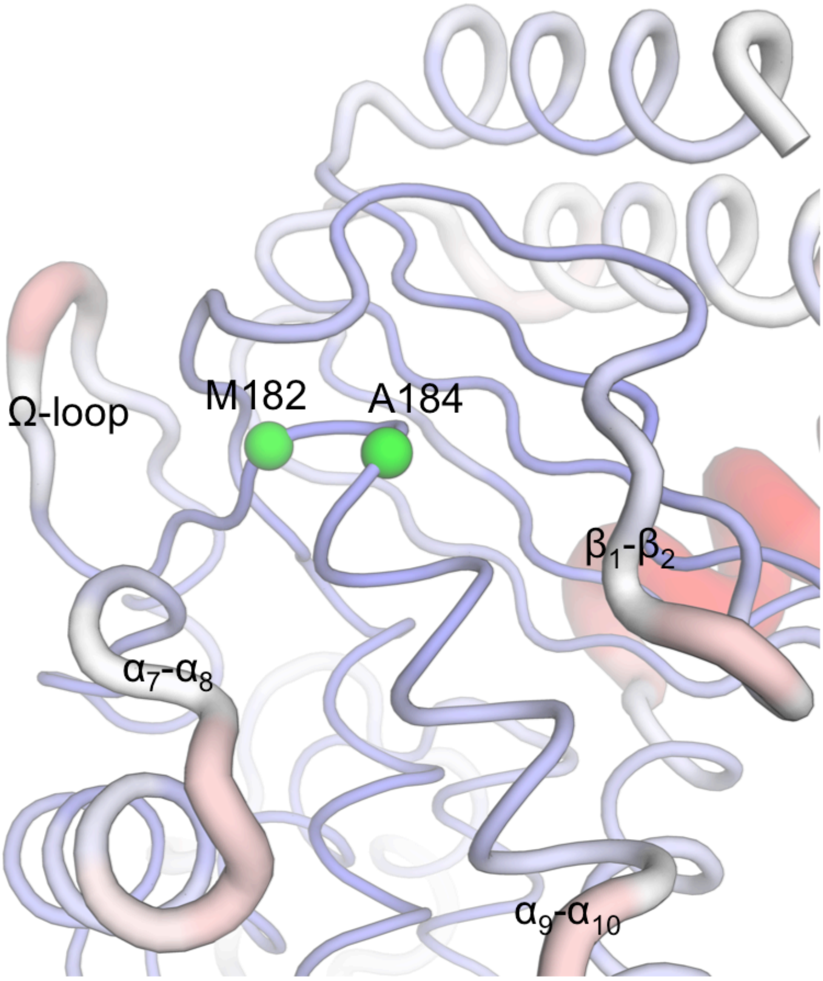


**Figure S16.** Spatial position of M182 and A184 on TEM-1. These residues (green spheres) are not on the communication pathway per se; but are in close vicinity and surrounded by the Ω loop, $\alpha_7$-$\alpha_8$ loop, $\beta_1$-$\beta_2$ loop and $\alpha_9$-$\alpha_{10}$ loop, that are involved in the communication network.

**Supplementary Movies:**

Signal propagation in TEM-1 and KPC-2 as a result of the perturbation (ligand removal) in the allosteric binding site. The disappearance of the ligand from its binding site generates a localized vacuum, against which there is an immediate structural response by the enzyme. As the simulation progresses, the cascading conformational changes in response to the perturbation (ligand removal) show the route by which structural response is transmitted through the protein.

**Movie S1**
Movie_S1-TEM1.mp4 (16 secs)
Signal propagation in TEM-1 (front view).

**Movie S2**
Movie_S2-TEM1.mp4 (16 secs)
Signal propagation in TEM-1 (back view).

**Movie S3**
Movie-S3-KPC2.mp4 (16 secs)
Signal propagation in KPC-2.

(14) Abreu, B.; Lopes, E. F.; Oliveira, A. S. F.; Soares, C. M. F508del Disturbs the Dynamics of the Nucleotide Binding Domains of CFTR before and after ATP

Hydrolysis. *Proteins Struct. Funct. Bioinforma.* **2020**, *88* (1), 113–126. https://doi.org/10.1002/prot.25776.
